# Kaposi’s Sarcoma-associated herpesvirus uses a novel protein fold to hijack RNA Polymerase II for viral late gene transcription

**DOI:** 10.64898/2026.06.29.734287

**Authors:** Pankaj Kumar Madheshiya, Shannon Henry, Xinyu Chen, Nicholas Doak, Taylor Gierke, Shaogeng Tang, Jimin Wang, Allison L. Didychuk

**Affiliations:** Department of Molecular Biophysics & Biochemistry, Yale University School of Medicine, New Haven, CT 06511, USA; Wu Tsai Institute, Yale University, New Haven, CT, USA

## Abstract

Herpesviruses, including the oncogenic human pathogen Kaposi’s Sarcoma-associated herpesvirus (KSHV), rely on cellular RNA Polymerase II (RNAPII) for expression of their protein-coding genes. Late in the lytic cycle, transcription initiation on a subset of KSHV genes depends on ORF24, a viral transcriptional activator that binds to viral late promoters through structural mimicry of a host transcription factor. Here, we present a structure of the N-terminal domain of ORF24, and show that it adopts a novel fold that facilitates direct binding of the RNAPII C-terminal domain (CTD). We further show that ORF24 is highly sensitive to the phosphorylation state of the RNAPII CTD, allowing for selective binding of the hypophosphorylated form to viral preinitiation complexes. By combining promoter binding and polymerase recruitment into a single protein, KSHV has streamlined essential steps in transcription complex assembly. Our work lays the groundwork for mechanistic dissection of a viral preinitiation complex.

---

Eukaryotic gene expression begins with binding of RNA Polymerase II (RNAPII) and general transcription factors to promoters, ultimately forming a preinitiation complex (PIC) composed of TFIIA, TFIIB, TFIID, TFIIE, TFIIF, TFIIH, and RNAPII.^3-6^ Canonical PIC assembly starts with recruitment of TFIID, whose central subunit TATA-binding protein (TBP) bends the DNA and interacts with TFIIA and TFIIB to initiate formation of the PIC.^7^ RNAPII is brought to the promoter in complex with TFIIF, followed by recruitment of TFIIE and TFIIH, promoter melting, and initiation of RNA synthesis.^8-13^ The human PIC also includes Mediator, a 26-subunit complex that helps position TFIIH and RNAPII and activate transcription initiation.^14-18^ Transcription initiation is a critical checkpoint in gene expression, determining the identity and abundance of resulting messenger RNAs.

Herpesviruses are large double-stranded DNA viruses that do not encode their own RNA polymerase and thus depend on cellular RNAPII for expression of viral protein-coding genes. The gammaherpesvirus Kaposi’s Sarcoma-associated herpesvirus (KSHV) is a driver of several human malignancies including Kaposi’s Sarcoma, primary effusion lymphoma, and multicentric Castleman’s disease. Neither prophylactic vaccines nor a cure for viral infection exist for any gammaherpesvirus. Therapeutic strategies largely focus on the treatment of KSHV-associated cancers, as many antiviral small-molecule drugs have limited efficacy^19^. Almost all anti-herpesvirus small molecules target a single step in the viral lifecycle – viral genome replication – including recently developed herpesvirus helicase/primase inhibitors.^20-22^

Like other herpesviruses, KSHV has a biphasic lifecycle, where replication of the viral genome and production of new infectious particles occurs during the lytic phase. Lytic genes are expressed in a tightly controlled temporal cascade, where replication of the viral genome distinguishes “early” and “late” genes.^23^ Transcription initiation from early viral promoters is thought to resemble the host PIC. By contrast, late gene expression in KSHV and related herpesviruses from the beta- and gammaherpesvirus subfamilies depends on six virus-encoded factors termed “viral transcriptional activators” (vTAs).^23^ One of these vTAs, encoded by ORF24 in KSHV, was identified as a viral mimic of cellular TBP.^24,25^ ORF24 functionally replaces TBP and uses its central TBP-like domain to directly bind the TATTWAA motif of the late gene promoter.^2,24-26^ In contrast to cellular TBP, which is incapable of stably recruiting RNAPII to promoters in the absence of other factors, ORF24 and its beta/gammaherpesvirus homologs were shown to interact with RNAPII by co-immunoprecipitation mass spectrometry.^24^ Thus, this viral TBP-like protein combines specific promoter binding and polymerase recruitment activities into a single protein.

We previously showed that the N-terminal domain (NTD) of ORF24, distinct from its TBP-like domain, is necessary and sufficient for interaction with RNAPII (**Fig. 1A**).^27^ The interaction site on RNAPII was identified as the unstructured C-terminal domain (CTD) of Rpb1, the largest subunit of RNAPII.^27^ In mammalian cells, the Rpb1 CTD consists of 52 copies of a repetitive heptad element with the consensus sequence Y1-S2-P3-T4-S5-P6-S7.^28^ The RNAPII CTD is a signaling hub that plays a crucial role in several stages of the transcription cycle, including initiation, elongation, co-transcriptional splicing, and termination.^29^ Post-translational modifications, particularly phosphorylation, are dynamically introduced throughout the transcription cycle, allowing for recruitment of distinct binding partners^30,31^. The unstructured CTD also drives clustering within nuclear condensates, presumably to form hubs that recruit and increase the local concentration of relevant processing factors *in vivo*.^32-34^

**Figure 1.**
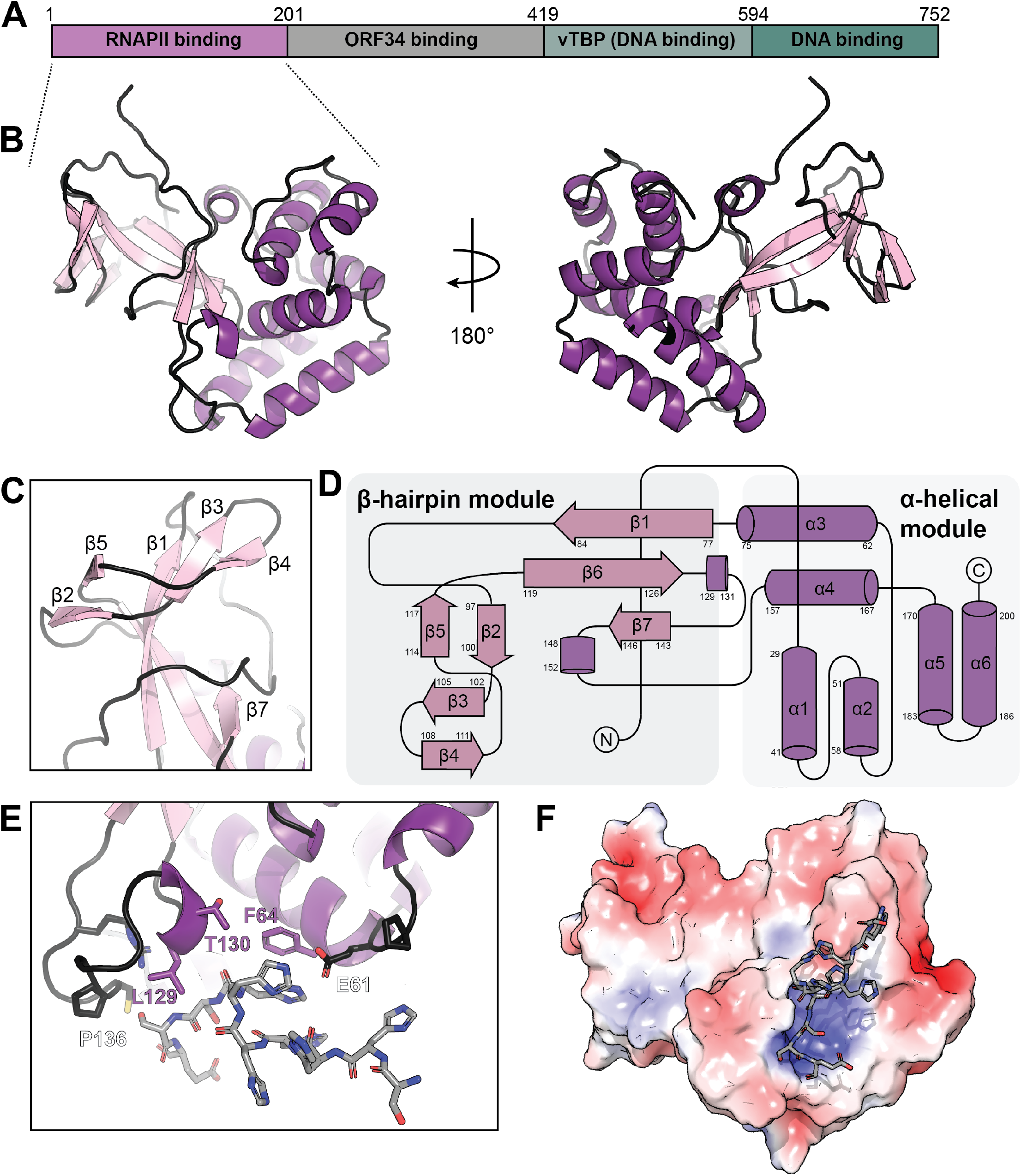
Structure of the N-terminal domain (NTD) of KSHV ORF24. **(A)** Domain organization and overall fold of ORF24-NTD. The N-terminal domain (residues 1-201) was used for structure determination. Residues 201-419 are implicated in binding to the vTA ORF34. Residues 419-594 are predicted to structurally resemble TATA-binding protein (TBP) and bind late gene promoters. Residues 594-752 have recently been shown to influence ORF24 DNA binding^2^. **(B)** Overall fold of ORF24-NTD with α-helices shown in purple and β-sheets in pink. **(C)** Close-up view of the β-hairpin. **(D)** Topological cartoon of ORF24-NTD with coloring as in (B). **(E)** Additional density attributed to the N-terminal hexahistidine purification tag was modeled (shown in grey sticks). **(F)** Electrostatic surface of ORF24-NTD, contoured from +5 kT/e (blue) to -5 kT/e (red).

ORF24 selectively immunoprecipitates hypophosphorylated RNAPII, consistent with a model where ORF24 and associated vTAs form a viral preinitiation complex.^24^ Here, we recapitulate selective recognition of unphosphorylated RNAPII CTD by ORF24 *in vitro* and show that this specificity is defined by the ORF24 NTD. To understand the basis of this specificity, we determined the structure of the ORF24-NTD and showed that it adopts a novel fold unique to herpesviruses. We identified surface-exposed residues that are required for interaction with RNAPII *in vitro* and during KSHV replication, laying the foundation for development of novel targeted therapeutics to disrupt this essential virus-host interaction.

## RESULTS

### The ORF24 polymerase recruitment domain adopts a novel fold

We first sought to structurally characterize the N-terminal domain (NTD) of ORF24. Residues 1-201 of ORF24 were recombinantly expressed and purified from *E. coli* with an N-terminal hexahistidine tag and TEV protease cleavage site. The protein was crystallized by vapor diffusion and crystals were obtained that diffracted X-rays to a maximum resolution of 1.93 Å (**Supplementary Table 1**). ORF24-NTD folds into a compact domain composed of a bundle of six α-helices interspersed with an extended β-hairpin (residues 77-126) (**Fig. 1B, C**). The α-helical bundle is well-ordered, whereas the β-hairpin and N-terminal loop exhibit higher B-factors. Residues 1-28 form an extended loop that wraps around the β-hairpin bundle (**Fig. 1D**). Dali^35^ and FoldSeek^36^ were used to identify potential structural homologs of ORF24-NTD outside the beta/gammaherpesviruses. Neither server aligned the domain to experimentally determined structures available from the Protein Data Bank, nor were there matches with high probability (for FoldSeek, probability > 0.5, E-Value < 1; for Dali, Z-score > 16 for a protein of 200 residues^37^). FoldSeek did identify a structural homolog within the AlphaFold Protein Structural Database (Uniprot A0A126LB08); however, this gene is misannotated as a human protein, and is actually the ORF24 homolog from the betaherpesvirus HHV-6A. Thus, ORF24-NTD adopts a novel fold that does not structurally resemble any known Pol II-interacting proteins or transcription factors, including domains of Mediator known to bind RNAPII CTD.^16,38^

Two cysteine residues (C67 and C176) are located on adjacent α-helices and are oriented ∼4 Å apart. We saw potential density for a disulfide bond in a lower-resolution dataset; however, we did not see this density in our higher-resolution dataset (**Fig. S1A**). Mutation to serine at position C67 has no effect on the ability of ORF24-NTD to interact with Rpb1 in a transient transfection co-immunoprecipitation assay, while mutation of C176 to serine eliminates binding (**Fig. S1B,C**). Thus, while a disulfide bond could form under some conditions and may influence heterogeneity within the crystal, it is unlikely that stabilization through this element is strictly required for efficient interaction with RNAPII.

Interestingly, we observed additional protein density in the crystal lattice that could not be attributed to ORF24-NTD. We modeled ten residues of a peptide nestled between helices 3 and 5 and a semi-structured loop (residues 128-141) with an occupancy of ∼0.60 (**Fig. 1E**). Potential stacking interactions between the peptide and ORF24 F64, along with polar interactions from residues ORF24 E61, R71, R143, and T130, contribute to binding along with hydrophobic contacts from L129 and P136. The peptide binds near an electrostatically positive pocket on the surface of ORF24-NTD created by R71 and R140, although it does not fully dock into this region (**Fig. 1F**). Given the high occupancy of the peptide in the crystal, we reasoned that the peptide likely comes from the N-terminal hexahistidine tag that was not removed prior to crystallization. The tag is insufficiently long to arise from the same molecule of ORF24-NTD, and instead likely stems from an adjacent asymmetric unit of ORF24 ∼13 Å away (**Fig. S1D**). Inclusion of the hexahistidine purification tag may have facilitated crystal packing, as we were unable to generate crystals when the tag was removed. To ensure that no other contaminating peptides were present, we performed ultra-performance liquid chromatography tandem mass spectrometry (UPLC-MS/MS) on both intact and desalted samples to assess the molecular weight and purity of our sample. The ORF24-NTD sample was highly pure and eluted at a molecular weight of 25,056 Da, slightly smaller than the predicted molecular weight of 25,188 Da, indicating that the initiating N-terminal methionine was removed (**Fig. S1E**). A tryptic digest followed by UPLC-MS/MS and search of the *E. coli* proteome for any peptides that may have co-purified during recombinant expression identified a single peptide from an *E. coli* recombinase present with comparable spectral counts to ORF24-NTD, with all others represented at least an order of magnitude less. Thus, the peptide present in our structure likely stems from the purification tag. Observation of this hexahistidine tag in our crystal structure implies that ORF24 is capable of binding and inducing partial order on a short flexible peptide.

### A conserved patch on ORF24-NTD mediates interaction with RNAPII CTD

We next leveraged our ORF24-NTD structure to identify a potential binding site for the RNAPII CTD. We identified an extended conserved patch spanning the face of the α-helical bundle (**Fig. 2A**). Interestingly, this conserved patch also contacts the putative N-terminal hexahistidine tag present in our crystal structure. We selected 15 residues across the surface of ORF24-NTD to test their contribution to the ORF24-RNAPII interaction. We transiently transfected plasmids encoding ORF24-NTD mutants into HEK293T cells and performed affinity purifications to quantify the fraction of Rpb1 bound for each mutant (**Fig. 2B, Fig. S2A**). Mutation of most of the highly conserved residues impacted binding, with a complete loss of binding when residues N42, F64, F65, R68, R71, T130, R143, R166, and E172 were mutated to alanine, whereas mutation of some highly conserved residues (E61, T63, R162) or less conserved positions (Q46, R95, Q175) to alanine minimally inhibited binding. When mapped onto the structure, residues that are critical for interaction with Rpb1 reside at the center of the conserved patch (**Fig. 2C**).

**Figure 2.**
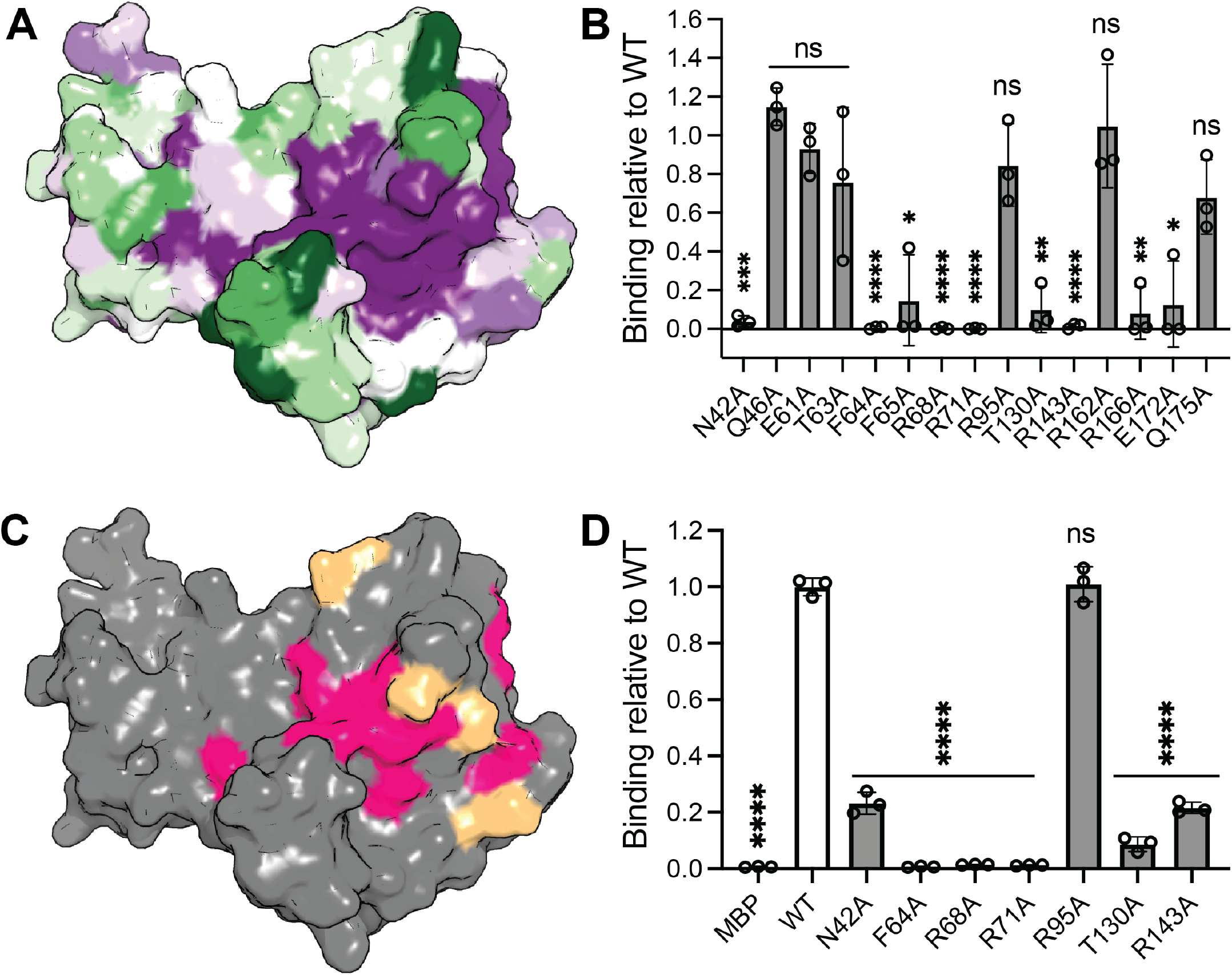
Identification of a conserved RNAPII binding site on ORF24-NTD. **(A)** Surface representation of ORF24-NTD colored by conservation, where purple residues are highly conserved and green residues are variable as determined by the ConSurf server ^1^. **(B)** Plasmids encoding wild-type or mutant strep-tagged ORF24-NTD were transiently transfected into HEK293T cells and subjected to affinity purification on StrepTactinXT beads followed by western blot analysis. Images were quantified and binding was normalized to WT ORF24-NTD in each experiment, with representative blots shown in **Supplementary Figure S2A**. Data are from three biological replicates, with statistics calculated using a one sample t-test where WT binding was set to 1; \*\*\*\**P* < 0.0001\*\*\**P* < 0.001 \*\**P* < 0.01 \*\**P* < 0.05. **(C)** Summary of co-immunoprecipitation data mapped on the surface of the ORF24-NTD structure. Residues determined to be important for the ORF24-NTD•RNAPII interaction are indicated in pink, while residues where mutation to alanine has minimal effect on RNAPII interaction are indicated in wheat. **(D)** Quantification of *in vitro* binding assay where recombinantly purified His-ORF24-NTD-strep wildtype and mutants were incubated with MBP-4xCTD followed by enrichment on StrepTactin XT beads, with representative gels shown in **Supplementary Figure S2B**. Data from three independent replicates were normalized to the average of wild-type binding, with statistics calculated using an unpaired t-test; \*\*\*\**P* < 0.0001.

To rigorously confirm that these mutations act directly on the ORF24-RNAPII interaction rather than by disrupting protein folding, we purified recombinant ORF24-NTD containing a representative subset of mutations and tested direct binding to the CTD *in vitro* using a purified construct with four CTD heptad repeats fused to MBP. Similar to what we saw in the cell-based experiments, ORF24 mutations F64A, R68A, R71A eliminated binding, mutations N42, T130A, and R143A greatly reduced the interaction, and mutation to R95A had no effect (**Fig. 2D, Fig. S2B**). All of the tested ORF24-NTD mutant proteins behaved similarly to WT ORF24-NTD during expression and purification, suggesting that mutations at these surface-exposed positions do not influence the fold of this isolated domain. In contrast, we were unable to purify ORF24-NTD harboring the L73A/L74A/L75A mutation that has previously been shown to disrupt the ORF24-Pol II interaction^24,26^ (**Fig. S2C**).

Residues 73-75 are buried within the structure, and mutation of these residues likely disrupts the fold of ORF24-NTD (**Fig. S2D**). Thus, we have identified single, surface-exposed residues that are key to the ORF24-RNAPII interface.

We noted that ORF24 residues 25 and 133 are variable between the KSHV BAC16 sequence (ultimately derived from the KSHV JSC1 sequence isolated from primary effusion lymphoma cells^39^) and our recombinant expression constructs, which match the sequence of ORF24 expressed in BCBL-1 cells (a different KSHV patient-derived line^40^). In BAC16, ORF24 residues 25/133 are aspartic acid [D]/tyrosine [Y], while in BCBL-1 cells and our expression constructs, these residues are tyrosine [Y]/cysteine [C]. Neither position is conserved across the gammaherpesviruses, even in the closely related rhadinovirus Colobine gammaherpesvirus 1 (**Fig. S3A**). Analysis of 270 ORF24 sequences deposited in GenBank revealed the presence of D/Y, Y/C, and D/C combinations at these positions, but none containing Y/Y (**Fig. S3B, Supplemental Table 2**). Residue 133 is immediately adjacent to the Rpb1 binding site (**Fig. S3C**) and is proximal to T130, whose mutation greatly reduces RNAPII binding (**Fig. 2**). Residue 25 is surface-exposed and located on the opposite face of the putative binding interface and is thus unlikely to contribute to the RNAPII interaction (**Fig. S3D**). We tested binding of all four combinations at these residues (Y/C, D/C, Y/Y, D/Y) in transient transfection co-immunoprecipitation experiments (**Fig. S3E,F**) and *in vitro* pulldowns (**Fig. S3G,H**). All combinations can bind to RNAPII, although the D/C combination appears to have a Interestingly, recombinant ORF24-NTD Y/Y expressed much better than other variants during purification, suggesting that this combination of residues could help folding or solubility. This combination does not appear to occur in circulating KSHV strains, raising the possibility that ORF24 expression or activity must be controlled for successful viral spread.

### The ORF24-NTD binding site is required for late gene expression and infectious virion production

To test the importance of the ORF24-RNAPII binding site in the context of viral infection, we introduced mutations N42A, F64A, R68A, and R95A into the ORF24 locus in the KSHV BAC16 genome using Red recombineering (**Fig. 3A**)^39^. ORF24 was endogenously tagged with an N-terminal HA epitope tag to facilitate detection^26^. We generated latently infected iSLK renal carcinoma cells harboring these recombinant viruses, allowing us to monitor progression through the viral lifecycle upon reactivation. We monitored viral gene expression after reactivation and observed that, consistent with our *in vitro* binding observations, the F64A and R68A mutant viruses exhibited a total loss of the representative viral late gene K8.1, the N42A mutation had an intermediate effect, and the R95A mutation had no effect (**Fig. 3B**). All mutant viruses expressed the representative viral early gene ORF6. We then tested whether these mutations inhibited production of infectious virus. Mutations that prevent interaction with Rpb1 (F64A and R68A) failed to generate any progeny virions, as late gene products are required for formation of viral capsids (**Fig. 3C**). The N42A mutation, which partially disrupts interaction with Rpb1 and incompletely disrupts late gene production, fully blocks infectious virion production, suggesting that a minimum threshold of late gene expression is required to produce infectious virus. Immunoprecipitation of ORF24 late in infection demonstrated that mutations N42A, F64A, and R68A eliminated interaction with Rpb1, while R95A had no effect (**Fig. 3D**). All mutations maintain an interaction with ORF34, its vTA binding partner. Mutations that abrogate binding to RNAPII exhibited lower ORF24 expression compared to WT or the R95A mutant. Interaction with RNAPII could help stabilize expression of ORF24, as recently proposed^2^. Thus, we have identified a critical region on ORF24 that is required for recruitment of RNAPII to late gene promoters and therefore required for infectious virion production.

**Figure 3.**
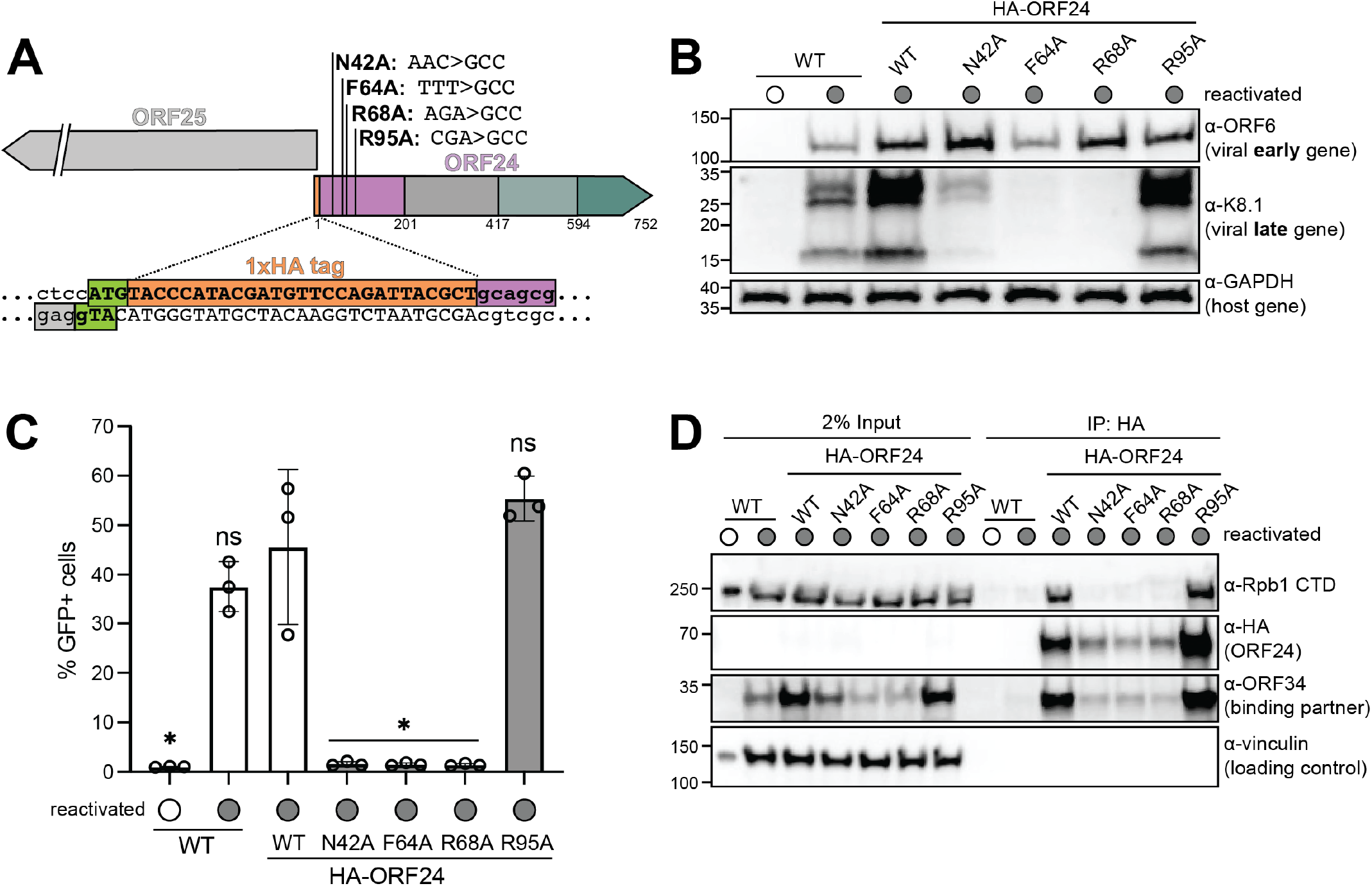
Mutations in the ORF24 RNAPII-binding interface block viral late gene expression. **(A)** Schematic representation of ORF24 locus in the KSHV BAC16 genome. An N-terminal HA epitope tag was introduced to facilitate protein detection. Residues N42, F64, R68, and R95 were mutated to alanine. **(B)** Western blot analysis of viral gene expression in iSLK cells infected with ORF24 mutant viruses. Where indicated, cells were reactivated for 72 hours. Data are representative of three biological replicates. **(C)** Infectious virion production from wild-type or variant ORF24-containing iSLK cells was monitored by supernatant transfer assay. GFP fluorescence of target HEK293T cells was measured via flow cytometry. Data are from three independent biological replicates, with statistics calculated using an unpaired t-test compared to HA-ORF24 WT; \**P* < 0.05. **(D)** Immunoprecipitations from unreactivated or reactivated wild-type, HA-tagged ORF24, or HA-tagged ORF24 harboring mutations at the indicated positions. Mutations at positions N42, F64, and R68 eliminate interaction with cellular RNAPII but maintain interaction with viral binding partner ORF34. Data are representative of three biological replicates. Molecular weight (in kDa) was determined via a protein ladder and is indicated to the left. Vinculin serves as a loading control.

### ORF24-NTD selectively recognizes hypophosphorylated RNAPII CTD

We next sought to understand how ORF24 binds the RNAPII CTD and selectively recruits hypophosphorylated Rpb1 CTD^24,27^. Previous work showed that full-length ORF24 fails to immunoprecipitate CTD harboring pS2, pS5, and pS7 modifications, but other states, including pY1 and pT4, were not tested^24^. We performed affinity purifications of ORF24-NTD or full-length ORF24 in HEK293T cells and probed with antibodies that recognize distinct phosphorylation states of the CTD. The Rpb1 CTD exists in two forms that exhibit different mobilities on an SDS-PAGE gel. The hypophosphorylated form (IIa) migrates at ∼250 kDa and allows for PIC formation, while the hyperphosphorylated form (IIo) migrates slower and is associated with active transcription^41,42^ Probing with the Rpb1 8WG16 antibody, which preferentially recognizes unphosphorylated heptads, we find that ORF24-NTD and full-length ORF24 bind hypophosphorylated Rpb1 to comparable extents, indicating that regions of ORF24 beyond the NTD likely do not contribute to RNAPII recruitment (**Fig. 4A**). Many commercial antibodies for phosphorylated CTD repeats recognize both IIo and IIa forms. Full-length and ORF24-NTD immunoprecipitated phosphorylated Rpb1 to similar extents using all of the antibodies tested; however, the resulting immunoprecipitated Rpb1 consistently migrated as IIa (**Fig. 4A**). Thus, ORF24 recognizes hypophosphorylated or minimally phosphorylated Pol II *in vivo*.

**Figure 4.**
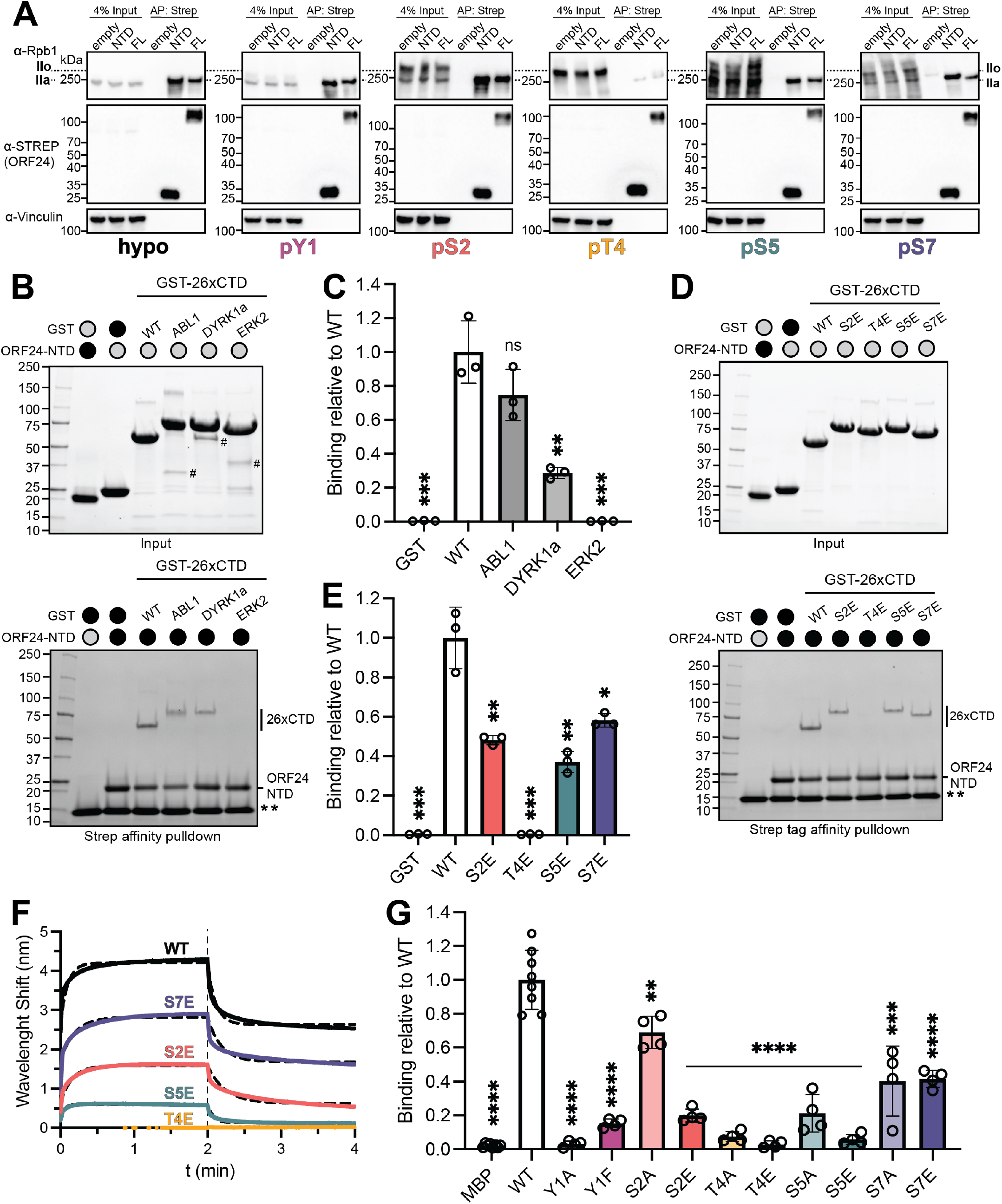
Mutations at all CTD heptad positions influence ORF24-NTD binding. **(A)** Plasmids encoding strep-tagged ORF24 full-length (FL) or ORF24-NTD were transiently transfected into HEK293T cells and subjected to affinity purification on StrepTactinXT beads followed by western blot analysis using antibodies specific for different phosphorylation states of the RNAPII CTD. ORF24 enriches the faster-migrating hypophosphorylated (IIa) state rather than hyperphosphorylated (IIo). Data are representative of three biological replicates. Molecular weight (in kDa) was determined via a protein ladder and indicated to the left. Vinculin serves as a loading control. **(B)** Representative *in vitro* binding experiment using phosphorylated RNAPII CTD. Phosphorylation of GST-26xCTD was performed with recombinantly purified ABL1, DYRK1a, or ERK2 kinases. Phosphorylated 26xCTD constructs were incubated with His-ORF24-NTD-strep and applied to StrepTactinXT beads. Free GST was used as a negative control. The top gel shows input (unphosphorylated and phosphorylated GST-26xCTD and His-ORF24-NTD-strep) while the bottom gel shows the resulting enrichment from StrepTactinXT beads. (#) indicates the kinase used for phosphorylation, (**) indicates a subunit of Strep-TactinXT released from the beads during boil elution. Gels were visualized by stain-free imaging and quantified in (C). **(C)** Three independent replicates of the phosphorylated GST-26xCTD binding experiment were quantified and normalized to wild-type (unphosphorylated GST-26xCTD) for each experiment. Statistics were calculated using an unpaired t-test; \*\*\**P* < 0.0005 \*\**P* < 0.005. **(D)** Representative *in vitro* binding experiment using phosphomimetic RNAPII CTD. GST-26xCTD constructs containing phosphomimetic mutants (S2E, T4E, S5E, and S7E in every heptad) were incubated with His-ORF24-NTD-strep and applied to StrepTactinXT beads. Free GST was used as a negative control. The top gel shows input (wild-type and mutant GST-26xCTD and His-ORF24-NTD-strep) while the bottom gel shows the resulting enrichment from StrepTactinXT beads. (**) indicates a subunit of Strep-TactinXT released from the beads during boil elution. Gels were visualized by stain-free imaging and quantified in (E). **(E)** Three independent replicates of the phosphomimetic GST-26xCTD binding experiment were quantified and normalized to wild-type GST-26xCTD for each experiment. Statistics were calculated using an unpaired t-test; \*\*\**P* < 0.0005 \*\**P* < 0.005 \**P* < 0.05. **(F)** Biolayer interferometry traces of ORF24-NTD binding to His-AviTag-MBP-26xCTD phosphomimetic mutants. Biotinylated His-AviTag-MBP-26xCTD variants (1 µM) were loaded on streptavidin sensors and binding to 48 µM His-ORF24-NTD was monitored. Data are representative of three independent replicates. **(G)** Mutations to all phosphorylatable positions in RNAPII CTD affect ORF24-NTD binding. An *in vitro* binding experiment was performed where MBP-4xCTD constructs containing mutations to Y1, S2, T4, S5, or S7 were incubated with His-ORF24-NTD-strep and applied to StrepTactinXT beads. Free MBP was used as a negative control. Representative gels are shown in **Supplementary Figure S4**. Four independent replicates of the mutant MBP-4xCTD binding experiment were quantified and normalized to wild-type MBP-4xCTD for each experiment. Statistics were calculated using an unpaired t-test; \*\*\*\**P* < 0.0001 \*\*\**P* < 0.001 \*\**P* < 0.01.

Given that we can recapitulate ORF24-CTD interactions using purified protein, we explored the effect of CTD phosphorylation by kinases implicated in transcription regulation. We performed *in vitro* binding assays using recombinantly purified ORF24-NTD and a CTD construct containing 26 heptad repeats fused to GST to more closely mimic the 52 repeats present in the human Rpb1 CTD. We used three diverse kinases to phosphorylate the 26xCTD construct. ABL1 is a non-receptor tyrosine kinase that can modify Y1.^43^ DYRK1a and ERK2 have been reported to modify both positions S2 and S5 *in vivo*, although DYRK1a is reportedly S2-selective while ERK2 is reportedly S5-selective *in vitro*.^44-46^ Phosphorylation of GST-26xCTD reduces its mobility on SDS-PAGE (**Fig. 4B**). To verify the presence of the appropriate modification, we blotted with phospho-specific antibodies (**Fig. S4A**). While ABL1 and ERK2 are specific for phosphorylation at Y1 and S5, respectively, in our conditions, DYRK1a phosphorylates both S2 and S5. Using the *in vitro* pulldown assay, we find that some ORF24 is capable of binding 26xCTD phosphorylated by ABL1 and DYRK1a, but not ERK2 (**Fig. 4B, C**). As a control, we confirmed that phosphorylation of GST-26xCTD does not result in nonspecific binding to the beads in the absence of ORF24 (**Fig. S4B**). These data suggest that introduction of negatively charged phosphate groups does not generally inhibit binding, and instead that ORF24 appears to depend on more specific recognition elements.

To define the sites and stoichiometry of mutations, we performed *in vitro* binding assays using 26xCTD constructs where every S2, T4, S5, or S7 residue was mutated to glutamic acid as a phosphomimetic and monitored ORF24 binding. Similar to the results with kinase-mediated phosphorylation, we find that ORF24-NTD is sensitive to mutations at all positions, but to differing extents (**Fig. 4D,E**). ORF24 can bind S2E, S5E, and S7E CTD constructs with reduced efficacy, and is least sensitive to the S7E mutation. In contrast, the T4E mutations eliminate binding. As a control, we confirmed that the 26xCTD phosphomimetic constructs do not bind the beads in the absence of ORF24 (**Fig. S4C**). To verify these binding patterns with an orthogonal method, we monitored interactions using biolayer interferometry (BLI), where different biotinylated 26xCTD constructs were loaded onto the sensor and ORF24 was recruited from solution (**Fig. 4F**). BLI appears to be more sensitive than pulldown experiments and allows for delineation of mutations that have similar effects, confirming that ORF24 is most affected by mutations at positions T4 and S5 and least affected by mutations at S7.

Since we can monitor ORF24-NTD binding to 4xCTD repeats, we introduced mutations to all five phosphorylatable residues (Y1, S2, T4, S5, and S7) and performed *in vitro* binding assays to more extensively explore ORF24 binding preferences. Serine and threonine residues were changed to either alanine or glutamic acid; Y1 was mutated to either alanine or phenylalanine. Alanine mutations were introduced to remove potential sidechain-specific interactions while glutamic acid was used as a mimetic for phosphorylated serine/threonine. All mutations to the CTD negatively affected ORF24-NTD binding (**Fig. 4G, Fig. S4D,E**). Mutation of either S2 or S7 to alanine had the smallest effect (30-60% loss of binding). Introduction of glutamic acid at position S2 had a larger effect than mutation to alanine, while S7 was affected to a similar extent by mutations to either alanine or glutamic acid. In contrast, ORF24-NTD binding was highly sensitive to either alanine or phosphomimetic mutations at positions T4 and S5, with phosphomimetics eliminating binding. ORF24-NTD was similarly sensitive to mutations at position Y1, as even the conservative Y1F mutation reduced binding ∼5-fold, with the Y1A mutation nearly eliminating binding. We confirmed that mutations in MBP-4xCTD do not result in nonspecific binding to the beads in the absence of ORF24 (**Fig. S4F**). Thus, ORF24-NTD is most sensitive to variations at sidechains Y1, T4, and S5 in the RNAPII CTD.

Several surprising findings emerge from these data. First, binding defects in the shorter 4xCTD construct can be alleviated with additional repeats (i.e., S5E in the context of 4xCTD vs. 26xCTD). Second, binding to phosphomimetic CTD does not perfectly correlate to binding after phosphorylation (i.e., 26xCTD S5E vs. ERK2-treated 26xCTD). Despite installation of both S2p and S5p by DYRK1a, ORF24-NTD can bind this substrate, whereas ERK2 activity (S5p modification alone) eliminates binding. Despite the clear reduction in mobility on a gel, it is possible that not all repeats are modified. Alternatively, phosphorylation at different sites may change the propensity of the intrinsically disordered CTD to adopt a peptide conformation that is disfavored for binding. Ultimately, the determinants of ORF24-NTD binding likely include sequence specificity, electrostatics, and backbone conformation that allow it to selectively recruit the appropriate phosphorylated state of RNAPII.

### ORF24 binds RNAPII with weak affinity and can be recruited to phase separated condensates

We previously demonstrated that ORF24 was capable of binding CTD constructs containing four or more heptapeptide repeats, but that two repeats were insufficient for binding^27^. Using BLI, we measured the affinity of ORF24 for the CTD and tested how the number of CTD heptad repeats influences ORF24-NTD binding. We recombinantly purified CTD constructs where different numbers of heptad repeats (0, 1, 2, 4, 10 and 26x) were fused to MBP; these were loaded onto the sensor with ORF24-NTD in solution. We confirmed that between two and four heptad repeats are minimally required to recruit ORF24-NTD, as binding was not detectable with 1x or 2x repeats (**Fig. 5A**). ORF24-NTD binds the CTD weakly with micromolar affinity. Longer repeats (10x, 26x) can recruit more copies of ORF24-NTD, resulting in a higher magnitude wavelength shift, but these constructs exhibit binding kinetics that are poorly fit by a simple association-dissociation nonlinear regression.

**Figure 5.**
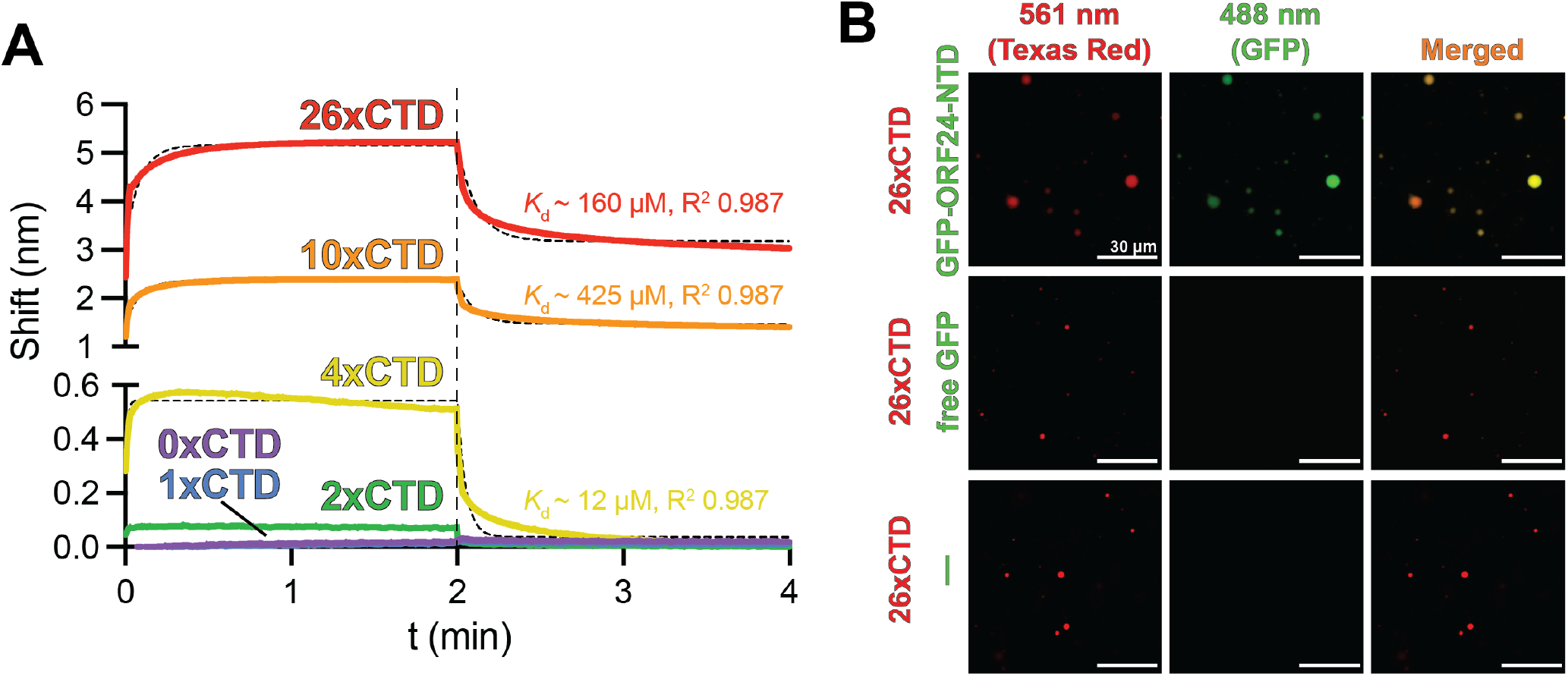
ORF24-NTD binds with CTD heptad repeats with weak affinity but can be incorporated into phase separated condensates. **(A)** Biolayer interferometric traces showing binding of ORF24-NTD to His-AviTag-MBP-xCTD fusions with either 0x (purple), 1x (blue), 2x (green), 4x (yellow), 10x (orange), or 26x (red) CTD consensus heptad repeats. Streptavidin sensors were loaded with 1 μM of biotinylated His-AviTag-MBP-xCTD and dipped into 48 μM of His-ORF24-NTD. Association and dissociation events were each monitored for 2 minutes. Curves were fit by nonlinear regression, represented by the black dotted line behind each curve. Data are representative of three independent replicates. Binding affinity (*K*_d_) and fit for each construct is indicated to the right. **(B)** ORF24-NTD is recruited to phase separated 26xCTD condensates. Free GFP and GFP-ORF24-NTD-strep do not form condensates alone. GFP-ORF24-NTD-strep, but not free GFP, is recruited into 26xCTD condensates. 10 µM of the indicated proteins were incubated in the presence of 16% dextran as a crowding agent prior to imaging. Scale bars indicate 30 µm.

While ORF24-NTD may directly recognize between 2-4 repeats, binding occurs in the context of the full-length (52x heptad repeat) RNAPII CTD, a low-complexity domain that can undergo phase separation. We used a previously established system where a recombinantly purified 26xCTD construct can be driven into phase separated condensates in the presence of crowding agents *in vitro*.^32,47^ We labeled GST-26xCTD with a fluorescent dye and observed condensation into droplets using confocal fluorescence microscopy (**Fig. 5B**). GFP-tagged ORF24-NTD is recruited to these CTD droplets, suggesting that it can bind RNAPII when it forms condensates. Free GFP is not recruited to CTD droplets, and addition of free GFP does not affect droplet formation. To test if ORF24-NTD is capable of undergoing phase separation, we used a turbidity assay, where increasing concentrations of protein are arrayed against increasing concentrations of the crowding agent dextran and absorbance at 350 nm is monitored. Notably, while GST-26xCTD phase separates at high dextran concentrations, ORF24-NTD does not undergo phase separation regardless of fusion to a GFP tag (**Fig. S5**). Thus, despite its relatively weak binding affinity for the RNAPII CTD, ORF24-NTD can be efficiently recruited into phase separated condensates even though this domain does not undergo phase separation itself.

## DISCUSSION

Here, we define how the oncogenic pathogen KSHV co-opts a core component of the eukaryotic transcription machinery to facilitate expression of critical viral genes at the end of its lifecycle. Our high-resolution structure of the RNAPII-interacting domain of the viral transcription factor ORF24 revealed a novel fold unique to herpesviruses. Multiple lines of evidence derived from conservation, mutation, and structural analysis allowed us to pinpoint a critical region on ORF24 that binds the RNAPII CTD. We showed that mutation of residues in ORF24 putatively involved in CTD binding prevented production of infectious virions due to a block at late gene expression. Given that ORF24-NTD does not structurally resemble human proteins and thus possesses a unique binding pocket, it may be possible to develop a new class of effective antiviral agents that target the essential ORF24-RNAPII interaction.

We used *in vitro* and *in vivo* binding assays to explore ORF24 selectivity for CTD phosphorylation state. The majority of mutations to the CTD reduced binding *in vitro*. Interestingly, some mutations (i.e., the phosphomimetic S7E), had a minimal (<2-fold) effect on binding *in vitro*, yet ORF24 was unable to immunoprecipitate pS7 RNAPII from human cells. Thus, while ORF24 effectively discriminates between most positions *in vitro*, selectivity is further enhanced *in vivo*. One potential explanation for this phenomenon is that the available pool of RNAPII in the cell exists at a concentration near the binding affinity of ORF24, and that selectivity is carefully tuned such that our *in vitro* equilibrium binding experiments at a single concentration do not fully capture binding behavior. Furthermore, RNAPII abundance and modification state likely change during infection. The total pool of RNAPII is decreased during KSHV lytic replication, including both pS2 and pS5 states^48^. In the related gammaherpesvirus MHV68, virus-driven host shutoff manifests as a loss of RNAPII at cellular genes caused in part by a reduction in nuclear RNAPII.^49^ Minor differences in *in vitro* binding affinity may result in different biological effects in the context of infection as the availability of RNAPII changes throughout the viral lifecycle.

An alternative hypothesis is that ORF24-NTD recognizes an untested modification of the CTD or nonconsensus heptad. Identification and interpretation of new post-translational modifications that comprise the CTD code is a growing and complex field.^50-53^ Noncanonical lysines at position 7 (CTD-K7) can be methylated and acetylated, and K7me1 and K7me2 modifications occur prior to transcription elongation^54^; monomethylated CTD-K7 migrates with hypophosphorylated CTD^55^, making it difficult to determine if ORF24 might interact with this modified heptad. Modification of noncanonical residues (R7me, K7ac) has been associated with transcription of specific gene classes.^56-58^ By analogy, ORF24 may have evolved to select a specifically modified CTD to facilitate viral late gene expression. It is also possible that prolines in the heptad sequence, which can undergo isomerization between *cis* and *trans* states and influence local structure, may play a role in selective recognition.^47,59^ Further investigation into the complex combinatorial landscape of human RNAPII sequence and modification will illuminate the precise binding determinants of ORF24.

ORF24 binds ∼2-4 hypophosphorylated consensus CTD heptad repeats with low micromolar binding affinity, consistent with the affinity of other CTD binding factors.^51^ Relatively weak affinity may be optimal, as the repetitive nature of the CTD and propensity to phase separate may drive multivalent interactions. ORF24 must achieve weak but selective binding, allowing it to compensate for the many repeats of the RNAPII CTD; dynamic binding would also allow it to sample multiple heptads. Binding in a cellular context is likely enhanced by avidity for multiple repeats and may be further fine-tuned by the physical restraints of the infected cell nucleus. Late in infection, synthesis of the viral genome results in “replication compartments” within the nucleus that dramatically reorganize the nucleus and change its physical properties.^60,61^ The newly replicated viral genome – the putative substrate for ORF24-mediated transcription initiation at viral promoters – is thought to lack classically defined chromatin.^62^ It is unclear how these changes influence the propensity of RNAPII to undergo phase-separation into transcriptional condensates, although transcription “factories” have been reported on KSHV genomes during lytic replication.^48^

Influenza A virus (IAV) also hijacks cellular RNAPII to facilitate viral gene expression.^63,64^ Both KSHV and IAV encode essential proteins that directly interact with the RNAPII CTD; however, they have evolved to bind different phosphorylation states of the CTD for distinct downstream uses. The PA subunit of IAV RNA-dependent RNA polymerase (FluPol) interacts with actively transcribing (pS5) RNAPII to perform “cap snatching” of host mRNA, while ORF24 recruits uninitiated (hypophosphorylated) RNAPII to viral promoters for transcription initiation.^64^ Although ORF24-NTD and PA are structurally unrelated, both domains require four heptad repeats for efficient binding.^65^ For FluPol, this requirement stems from the presence of two CTD binding sites that each recognize ∼1 heptad repeat and supply positively charged binding pockets that confer specificity for pS5.^65-67^ ORF24 exhibits weaker binding for its preferred CTD substrate than FluPol (∼12 μM vs. ∼1 μM).^65^ The improved avidity stemming from two distinct binding sites in FluPol likely contributes to its higher affinity, although other contacts between FluPol and RNAPII may further improve binding.^68,69^ It is formally possible that other elements within ORF24 or its associated vTAs contribute to binding to other elements within Rpb1 or RNAPII. However, our observation that full-length ORF24 and its NTD immunoprecipitate comparable amounts of Rpb1 from cells implies that the ORF24-NTD is the primary domain involved in RNAPII recruitment to late KSHV promoters.

Our data, particularly the selective recognition of hypophosphorylated RNAPII CTD, are consistent with a model of late gene transcription where viral factors form a unique viral preinitiation complex. Chromatin immunoprecipitation during viral infection showed that ORF24 and its associated viral transcription factor binding partners form a complex on viral late gene promoters.^24,26,70^ ORF24, in addition to its N-terminal polymerase recruitment domain, contains an additional domain predicted to structurally resemble the cellular transcription factor TBP, with specificity for a noncanonical TATT motif.^2,24,25^ While TFIID can promote transcription and bind TATA-less promoters through contact of TBP-associated factors (TAFs) to downstream regions^71,72^, binding of ORF24 to the TATT promoter element may preclude binding of TFIID.

We used AlphaFold 3^73^ to predict the structure of ORF24 in complex with the other KSHV vTAs (ORFs 18, 30, 31, 34, and 66) on a late gene promoter (**Fig. 6A**). We aligned the vTBP-like domain of ORF24 with TBP in the structure of the human Mediator-bound PIC (**Fig. 6B**).^16^ Binding of the other vTAs, particularly ORF18, 31, and 66, likely sterically occludes binding of general transcription factors TFIIA, TFIIB, and TFIIF, suggesting that promoter recognition by the vTAs is fundamentally altered despite the use of a TBP-like protein (**Fig. 6C**). It is unclear whether TFIID is required for viral late gene transcription, given that TFIID centers around TBP. However, TAF-containing complexes lacking TBP have been reported, raising the possibility that these factors are co-opted during viral infection.^72,74^

**Figure 6.**
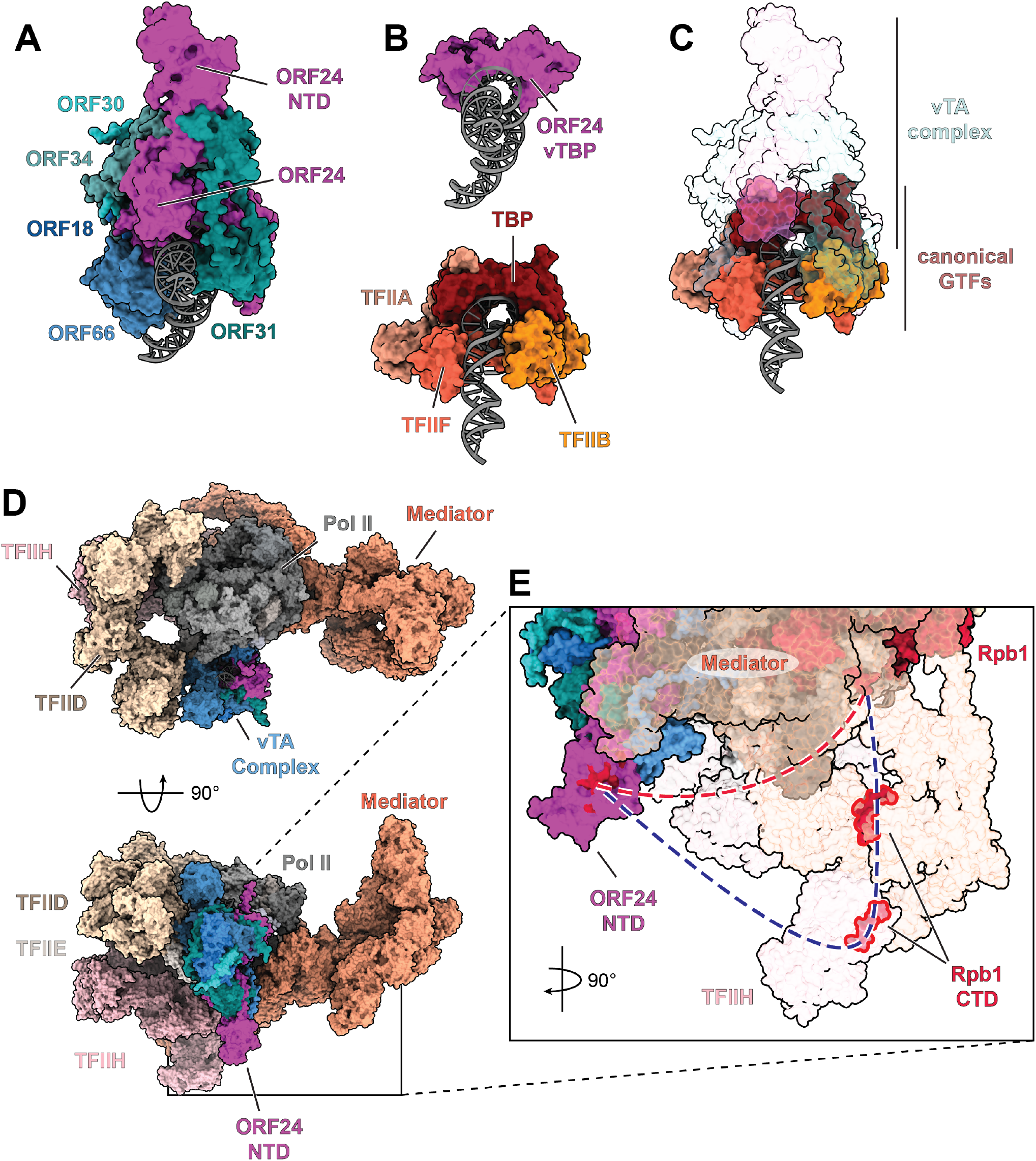
Model of the KSHV vPIC. **(A)** AlphaFold 3 model of the KSHV vTA complex (ORF18 in dark blue, ORF24 in magenta, ORF30 in dark turqouise, ORF31 in dark cyan, ORF34 in cadet blue, and ORF66 in steel blue) bound to the KSHV K8.1 promoter (grey). The vTA complex prediction was overlaid with the structure of the human Mediator-bound transcription pre-initiation complex (PDB 7LBM) based on structural homology between the ORF24 vTBP-like domain and human TBP. **(B)** Top: AlphaFold 3 prediction of the viral TBP-like domain of ORF24 (residues 419-594) bound to the KSHV K8.1 promoter in the context of the vTA complex. Bottom: selected general transcription factors TFIIA (dark salmon), TFIIB (dark orange), TFIIF subunit 2 (tomato) in the context of the human Mediator-bound PIC (PDB 7LBM) on the super core promoter (grey). **(C)** Overlay of the vTA complex prediction (transparent) on selected human general transcription factors (solid). **(D)** Human Mediator-bound PIC (PDB 7LBM; RNA Polymerase II in shades of grey, Mediator in light salmon, TFIIE in seashell, TFIIH in pink) with overlaid TFIID (peach) bound to promoter DNA (PDB 6MZM) and vTA complex prediction (colors as in (A)). TFIID was overlaid by alignment of promoter DNA to PDB 7LBM while the vTA complex was overlaid based on structural homology between the ORF24 vTBP domain and human TBP in 7LBM. **(E)** Potential paths of Rpb1 CTD when bound to the ORF24-NTD (magenta, with regions important for the interaction highlighted in red) independently (red dashed line) or concomitantly with Mediator and TFIIH (blue dashed line).

While the vTA complex is unlikely to co-exist as predicted with canonically-bound TFIIA and TFIIB, the orientation of the predicted complex suggests that it would not prevent binding of Mediator or TFIIH (**Fig. 6D**). Indeed, experimental data indicate that TFIIH is still required for herpesvirus late gene transcription.^24,75^ Mediator helps display the CTD to TFIIH for phosphorylation, triggering escape from the PIC.^16,18^ Binding sites for several CTD repeats in the PIC have been resolved, suggesting the path of the CTD (**Fig. 6E**).^16^ Given the length of the Rpb1 CTD, ORF24-NTD could bind a distal CTD repeat concomitant with Mediator/TFIIH binding (**Fig. 6E**). ORF24-NTD may enhance recruitment of Mediator, as proposed for the alphaherpesvirus transcription factor ICP4.^76^

Alternatively, ORF24-NTD could functionally replace multiple general transcription factors by eliminating the need for Mediator through its ability to bind the RNAPII CTD. If Mediator or TFIID (∼1.4 and 1.3 MDa complexes, respectively) are not required for late gene transcription, viral promoter complexes must be drastically simplified compared to the canonical ∼4 MDa human PIC. Consistent with this hypothesis, high-resolution chromatin immunoprecipitation sequencing in a related betaherpesvirus indicates that the vPIC has a smaller binding footprint on the promoter than a TBP-centered PIC.^77^ Further mechanistic dissection of this viral complex may reveal additional examples of viral structural innovation and illuminate the minimal components required to drive efficient transcription initiation. Understanding how ORF24 and its binding partners influence the assembly of preinitiation complexes in the context of viral infection will require integration of information from atomic to cellular scales.

## MATERIALS AND METHODS

### Cloning and expression constructs

ORF24-NTD, GFP, GFP-ORF24-NTD and CTD repeats (wild type and mutants) were cloned into pET28b or pQLink bacterial expression plasmids using ligation-independent cloning. Plasmids encoding GST-tagged 26xCTD (wild type, T4E, S5E, and S7E) and DYRK1a and ERK2 kinases were gifted by Y. Jessie Zhang. A geneblock encoding 26xCTD S2E was synthesized (Twist Bioscience) and cloned into the pET28a-GST backbone using ligation independent cloning. Plasmids for transient transfection of C-terminal 2xStrep-tagged ORF24-NTD or full-length ORF24 were previously described (Addgene #138442, #129742). Mutations in these plasmids were introduced by inverse PCR mutagenesis. Details for all plasmids are provided in **Supplementary Table 3** and all plasmids created in this study have been deposited in Addgene (#257637-257693).

### Protein expression and purification

#### ORF24 and GFP constructs

His-tagged ORF24-NTD, ORF24-NTD^-^strep, GFP-ORF24-NTD^-^strep and GFP proteins were expressed in NiCo21 (DE3) *E. coli* (New England Biolabs). Cells were grown in autoinduction media ^78^ at 37° C until the culture reached an OD_600_ ∼0.8-1.0, then the temperature was reduced to 18° C and the culture was grown for 16 hours. Cultures were harvested by centrifugation at 5,000 × g for 10 min at 4°C. The cell pellet was either frozen or immediately resuspended in lysis buffer (500 mM NaCl, 50 mM HEPES acid, 50 HEPES base, 1 mM TCEP, 20 mM imidazole, 0.1% Triton X-100 (except for GFP prep), 10% glycerol, 1 mM PMSF, pH 7.5) and lysed through sonication. Cell lysates were cleared by centrifugation at 40,000 × g for 1 hour at 4°C. Clarified lysate was incubated with Ni^2+^ agarose resin (GoldBio) for 45 minutes with rotation at 4°C. After binding, the resin was washed with fifty column volumes (CV) of wash buffer (500 mM NaCl, 50 mM HEPES acid, 50 HEPES base, 1 mM TCEP, 50 mM imidazole, 10% glycerol, pH 7.5), then bound fractions were eluted with five CV of elution buffer (500 mM NaCl, 50 mM HEPES acid, 50 HEPES base, 1 mM TCEP, 300 mM imidazole, 10% glycerol, pH 7.5). Elution fractions were concentrated using a 10 kDa centrifugal concentrator (MilliporeSigma) and loaded onto a Superdex 200 HiLoad 16/600 column or Increase 10/300 GL (Cytiva) pre-equilibrated in sizing buffer (100 mM NaCl, 20 mM HEPES acid, 20 HEPES base, 1 mM TCEP, 5% glycerol, pH 7.5). Proteins were assessed for purity using SDS-PAGE and visualized with Coomassie Brilliant Blue staining prior to aliquoting and storage at -80°C.

#### MBP-4xCTD used in pulldown experiments

Wild-type and mutant MBP-4xCTD protein was expressed as above. After harvesting, the culture pellet was either frozen or immediately resuspended in lysis buffer (300 mM NaCl, 50 mM HEPES acid, 50 HEPES base, 5 mM DTT, 5% glycerol, 1 mM PMSF, pH 7.4) and lysed through sonication. Cell lysates were cleared by centrifugation at 40,000 × g for 1 hour at 4°C. Clarified lysate was incubated with amylose agarose beads (New England Biolabs) for 2 hours with rotation at 4°C. After binding, the resin was washed with fifty CV of wash buffer (300 mM NaCl, 50 mM HEPES acid, 50 HEPES base, 5 mM DTT, and 5% glycerol, pH 7.4), then bound fractions were eluted with five CV of elution buffer (300 mM NaCl, 50 mM HEPES acid, 50 HEPES base, 5 mM DTT, 10 mM maltose, 5% glycerol, pH 7.4).

#### GST-26xCTD used in pulldown experiments

Wild-type 26xCTD and phosphomimetic mutants were expressed as above, except that cells were grown for 20 hours at 16°C. The cell pellet was either frozen or immediately resuspended in the lysis buffer (500 mM NaCl, 50 mM HEPES acid, 50 HEPES base, 5 mM DTT, 10% glycerol, 0.1% Triton X-100, 1 mM PMSF, pH 7.4) and lysed through sonication. Cell lysates were cleared by centrifugation at 40,000 × g for 1 hour at 4°C. Clarified lysate was incubated with glutathione agarose beads (Pierce) for 4 hours with rotation at 4°C. After binding, the resin was washed with fifty CV of wash buffer (500 mM NaCl, 50 mM HEPES acid, 50 mM HEPES base, 10% glycerol, 5 mM DTT, pH 7.4) and bound fractions were eluted with five CV of elution buffer (500 mM NaCl, 50 mM HEPES acid, 50 mM HEPES base, 10% glycerol, 5 mM DTT, 10 mM reduce glutathione, pH 7.4). Elution fractions were concentrated using a 10 kDa centrifugal concentrator (Millipore) and loaded onto a Superdex 200 HiLoad 16/600 column (Cytiva) in sizing buffer (300 mM NaCl, 50 mM HEPES acid, 50 mM HEPES base, 5% glycerol, 5 mM DTT, pH 7.4).

#### CTD variants used in biolayer interferometry analysis

His-AviTag-MBP tagged xCTD (variants and mutant protein) were expressed in CVB101 cells (Avidity LLC), which harbor an IPTG-inducible copy of the biotin ligase BirA. Cells were grown in Luria-Bertani (LB) medium at 37°C until the OD_600_ reached between 0.8-1.0, then biotin (Sigma-Aldrich; 50 µM final concentration) was added into the culture and the temperature was reduced to 18°C. After 1 hour of incubation, the culture was induced with 300 µM IPTG for 20 hours. Cultures were harvested by centrifugation at 5,000 × g for 10 min at 4°C. The culture pellet was either frozen or immediately resuspended in lysis buffer (300 mM NaCl, 50 mM HEPES acid, 50 mM HEPES base, 5% glycerol, 1 mM TCEP, 20 mM imidazole, 1 mM PMSF, pH 7.4) and lysed through sonication. Cell lysates were clarified by centrifugation at 40,000 × g for 1 hour at 4°C. Clarified lysate was incubated with Ni^+2^ agarose beads (GoldBio) for 45 minutes with rotation at 4°C. After binding, the resin was washed with fifty CV of wash buffer (300 mM NaCl, 50 mM HEPES acid, 50 mM HEPES base, 5% glycerol, 1 mM TCEP, 50 mM imidazole, pH 7.4) and bound fractions were eluted with five CV of elution buffer (300 mM NaCl, 50 mM HEPES acid, 50 mM HEPES base, 5% glycerol, 1 mM TCEP, 400 mM Imidazole, pH 7.4). Elution fractions were concentrated using a 10 kDa centrifugal concentrator (Millipore) and loaded onto a Superdex 200 HiLoad16/600 column (Cytiva) equilibrated in sizing buffer (300 mM NaCl, 50 mM HEPES acid, 50 mM HEPES base, 5% glycerol, 1 mM TCEP, pH 7.4). To ensure that proteins were biotinylated, 10 µg of purified protein was mixed with 2.5 µg of streptavidin and incubated for 1 hour at 25°C, then binding was quantified by a SDS-PAGE gel shift assay.

#### Purification of kinases for in vitro phosphorylation

His-tagged ERK2 and His-GST-tagged DYRK1a kinases were expressed in NiCo21 (DE3) *E. coli* cells using the same expression and purification conditions described for the nonbiotinylated CTD variants. Kinases were purified using Ni^+2^ agarose affinity pull-down followed by size exclusion chromatography. Recombinant human His-tagged ABL1 kinase domain (residues 248-531) was purchased from BPS Bioscience.

#### CTD *in vitro* phosphorylation

Phosphorylation of purified GST-26xCTD was performed based on previous work ^47,79^. In brief, for ERK2 or DYRK1a phosphorylation reactions, 170 µg of GST-26xCTD was incubated with 0.125 mg/mL ERK2 or DYRK1a in a 150 µL reaction containing 300 mM NaCl, 50 mM HEPES acid, 50 mM HEPES base, 5% glycerol, 1mM TCEP, 5 mM ATP and 5 mM MgCl2. For Abl1 phosphorylation reactions, 100 µg of GST-26xGST was incubated with 0.02 mg/mL Abl1 in 100 µL reaction containing 100 mM NaCl, 25 mM Tris pH 7.25, 5 mM MgCl_2_, 5 mM MnCl_2_, 1 mM DTT, 5% glycerol, 100 ng/µL BSA, 1 mM sodium orthovanadate and 5 mM ATP. Reactions were incubated at 30°C for 16 hours in a thermocycler. After incubation, phosphorylation was quenched via addition of 5 mM EDTA and quantified by SDS-PAGE gel shift assay.

### Crystallization of ORF24-NTD and structure determination

Crystals of His-ORF24-NTD were obtained by sitting drop vapor diffusion by mixing 0.2 µL of 5.6 mg/mL protein in sizing buffer (100 mM NaCl, 20 mM HEPES acid, 20 HEPES base, 1 mM TCEP, and 5% glycerol at pH 7.5) with 0.2 µL of reservoir buffer (0.15 M DL-Malic acid pH 7.0, 20% w/v polyethylene glycol 3350, 110 mM pyruvic acid). Crystal growth was observed after three days. A single crystal was looped into cryoprotectant solution (reservoir buffer supplemented with 25% glycerol) and directly frozen in liquid nitrogen. Diffraction data from a single crystal was collected on beamline BL12-2 of the Stanford Synchrotron Radiation Lightsource (SSRL). Data were integrated using XDS and scaled in XSCALE ^80^. Phases were determined by molecular replacement in PHASER using an AlphaFold 3 prediction of ORF24-NTD as a search model ^73^. Model building was performed in Coot ^81^ with refinement in Refmac5 as implemented in CCP4 ^82,83^.

#### *In vitro* pulldown assays

To quantify the interaction between ORF24-NTD and various CTD constructs (wild-type, mutant, or *in vitro* phosphorylated GST-26xCTD), 10 μg of recombinantly purified His-ORF24-NTD-Strep was mixed with 20 µg recombinantly purified CTD construct and 10 µL of pre-equilibrated MagStrep “type 3” XT beads (IBA Lifesciences). The final volume was adjusted to 300 µL using reconstitution or binding buffer (100 mM NaCl, 20 mM HEPES acid, 20 HEPES base, 1 mM TCEP, 5% glycerol, 0.05% NP-40, pH 7.4). Binding was performed with constant rotation at 4°C for 1 hour. The beads were washed three times for 5 minutes each and bound fractions were eluted by boiling in 2x Laemmli protein loading dye followed by SDS-PAGE analysis. At least three independent experiments were performed, the protein bands were quantified by ImageJ and analysis was performed using GraphPad Prism 10.14.0.

### Biolayer Interferometry (BLI) analysis

BLI experiments were performed on an Octet R8 system (Sartorius) using 96-well flat-bottom black plates (Greiner Bio-One). Streptavidin (SA) biosensor tips (Sartorius Octet SA biosensors) were pre-equilibrated in Octet buffer (100 mM NaCl, 20 mM HEPES acid, 20 mM HEPES base, 5% glycerol, 1 mM TCEP, 0.1% bovine serum albumin and 0.05% Tween20, pH 7.4) for 20 minutes. After equilibration, SA biosensor tips were loaded with 1 µM of biotinylated wild-type, mutant, or phosphorylated His-AviTag-MBP-CTD, dipped into 48 µM His-ORF24-NTD analyte for the association step, then returned to the baseline wells for the dissociation step. Binding curves were fit using Octet Analysis Studio 13.0 software using a 1:1 binding model, with background subtraction using a His-ORF24-NTD loaded sensor tip dipped into sample wells containing only Octet buffer. At least three independent experiments were performed. Analysis was performed using GraphPad Prism 10.14.0.

### Untargeted, label-free proteomic analysis following proteolysis via Ultra-High Performance Liquid Chromatography Coupled to Tandem Mass Spectrometry (UPLC-MS/MS)

To robustly identify and quantify ORF24-NTD and contaminating peptides/proteins in the purified solution, an aliquot containing 35 µg of purified protein was removed and subjected to bottom-up proteomic analysis via UPLC-MS/MS. Briefly, the purified proteins were diluted with 0.1 M ammonium bicarbonate in water and subjected to Cys reduction and alkylation at 25°C using a 1.5 hour incubation in 5 mM dithiothreitol followed by a 45 minutes incubation with 10 mM iodoacetamide in the dark. Carbamidomethylated proteins were proteolyzed using Promega modified sequencing grade trypsin at an enzyme:protein ratio of 1:20 for 16 hours at 37°C. After quenching the digestion using formic acid, the resulting peptides were desalted using Pierce peptide desalting spin columns using manufacturer’s instruction. The eluted peptides were dried to completion in a Labconco speedvac concentrator and stored at -20°C prior to analysis.

Just prior to untargeted UPLC-MS/MS analysis, the desalted peptides were thawed and resuspended in 0.1% formic acid in water (Solvent A) and quantified via A280 absorbance measurements on a Nanodrop spectrophotometer (Thermo Scientific). An aliquot of the resuspended peptides were diluted using Solvent A to 0.25 mg/mL and an aliquot containing 250 ng was loaded onto a nanoEase m/z Peptide BEH C18 analytical column (1.7 μm, 75 μm × 250 mm, Waters Corporation) using a Thermo Scientific Ultimate 3000 RSLCnano UPLC system and separated using a 300 nL/minute reversed phase UPLC gradient utilizing Solvent A and Solvent B (0.1% formic acid in acetonitrile) as follows: 10 minutes hold at 4% Solvent B, ramp from 4% Solvent B to 30% Solvent B in 50 minutes, followed by a ramp from 30% to 90% Solvent B in 10 minutes. Peptides were eluted directly into an Orbitrap Eclipse Tribrid mass spectrometer (Thermo Scientific) implementing a TopN data-dependent acquisition method for MS/MS analysis using the following parameters: capillary voltage of 2200 V, MS1 scan resolution of 120,000 and scan range of 300 – 1,800 m/z, MS2 scans acquired for charge states +2 to +8, MS2 isolation window of 1.6 m/z, a TopN cycle time of 3 s, and dynamic exclusion turned “on” for 30 seconds per precursor fragmented using a 20 ppm window.

Peptide and protein identifications and label-free quantification were achieved using MaxQuantREF (v2.0.2.0) ^84^ by searching against the HT-ORF24NTD sequence plus the Uniprot *E. coli* BL21(DE3) reference proteome (UP000002032, accessed 01/02/2025) with the following parameters: enzyme specificity: trypsin/P with up to 2 missed cleavages, minimum peptide length of 5, variable modifications of acetylation of protein N-terminus, deamidation of Asn/Gln, peptide N-terminal Gln to pyro-Glu conversion, and oxidation of Met, plus carbamidomethyl Cys set as a fixed modification. False discovery rates at the peptide-spectrum-match and protein levels were set to 1%. All other parameters were kept at default values. MaxQuant output was uploaded into Scaffold (v5, Proteome Software) for further analysis and data visualization.

### Intact His-ORF24 protein and peptide analysis via UPLC-MS and UPLC-MS/MS

To confirm the molecular weight of purified ORF24-NTD and search for abundant non-ORF24-NTD peptides present in the purified solution, two intact protein UPLC-MS experiments were conducted: once with sample clean-up using Pierce peptide desalting spin columns using manufacturer’s instruction, and once without clean-up to prove no contaminant peptide losses during desalting. For the experiment without desalting, a 15 pmol injection of the purified solution was loaded onto a nanoEase m/z Peptide BEH C4 analytical column (1.7 μm, 75 μm × 100 mm, Waters Corporation) using a Thermo Scientific Vanquish Flex UPLC system and separated using an identical flow rate and 1 hour binary gradient to that used for bottom-up proteomic analysis described above. Ions were eluted directly into a Q Exactive HF mass spectrometer (Thermo Scientific) operating with the following acquisition parameters: capillary voltage of 1.8 kV, MS1 resolution of 60,000, AGC target of 1E6, MS1 scan range of 300 to 1,800 m/z, Top3 MS/MS data-dependent acquisition, MS2 resolution of 15,000, AGC target of 1E5, isolation window of 2.0 m/z, charge exclusion “on” for unassigned and >+4 charge states, plus an exclusion duration of 15 seconds. Protein/peptide signal observed in the MS1 spectra was deconvolved using UniDecREF ^85^ to calculate the neutral protein molecular weight.

For the experiment that incorporated desalting, a 100 ng aliquot was loaded onto a nanoEase m/z Peptide BEH C18 analytical column (1.7 μm, 75 μm × 250 mm, Waters Corporation) using a Thermo Scientific Vanquish Neo UPLC system and separated using the same 1 hour binary reversed phase gradient. Ions were eluted directly Ions were eluted directly into a Q Exactive HF mass spectrometer (Thermo Scientific) operating with the same acquisition parameters as the first intact experiment except a Top15 data-dependent acquisition for MS2 was used. The resulting UPLC-MS/MS data were searched using MaxQuant (v2.0.2.0) using the same parameters described above.

### Tissue culture

HEK293T cells (ATCC CRL-3216) were maintained in Dulbecco’s Modified Eagle Medium (DMEM) supplemented with 10% fetal bovine serum (Gibco) at 37°C in 5% CO_2_ (v/v). HEK293T cells expressing N-terminally 1xHA epitope-tagged ORF24 (HEK293T-ORF24) were maintained in DMEM+FBS with 500 μg/mL zeocin. iSLK-puro cells were maintained in DMEM+10% FBS with 1 μg/mL puromycin. After establishment of latent infection with KSHV encoded on BAC16, iSLK-BAC16 cells were cultured in DMEM+10% FBS with 1 μg/mL puromycin and 1 mg/mL hygromycin. Cell lines were tested with a Mycoplasma Pro PCR Detection Kit (Applied Biological Materials Inc.).

### Immunoprecipitation

Transient transfection of plasmids expressing ORF24 (full-length, NTD, and mutants) into HEK293T cells was performed in a 15 cm dish at ∼70% confluency with 32 μg plasmid and 80 μg PEI MAX (MW 40,000) transfection reagent (Polysciences, Inc.); cells were harvested 24 hours post-transfection. For experiments using infected cells, 2.6 × 10^6^ iSLK cells were seeded in 15-cm dishes and reactivated the next day using 5 μg/mL doxycycline and 1 mM sodium butyrate; cells were harvested 72 hours post-reactivation. Cells were lysed in lysis buffer (150 mM NaCl, 50 mM Tris pH 7.4, 0.5% NP-40) supplemented with EDTA-free protease inhibitor cocktail (Roche) with rotation at 4°C for 30 minutes. Lysates were clarified by centrifugation at 21,000 × g for 10 minutes at 4°C. Total protein concentration was determined by Bradford assay, and 1 mg of lysate was incubated with 20 μL of pre-washed MagStrep Strep-TactinXT beads (IBA Lifesciences) in 1 mL of IP buffer (150 mM NaCl, 50 mM Tris pH 7.4) supplemented with EDTA-free protease inhibitor cocktail (Roche). After rotating at 4°C for 16 hours, the beads were washed three times in wash buffer (150 mM NaCl, 50 mM Tris pH 7.4, 0.05% NP-40). Protein was eluted by boiling in 2x Laemmli Sample Buffer (Bio-Rad) supplemented with β-mercaptoethanol. Input samples (2% of total IP - 20 μg) were incubated with 4x Laemmli Sample Buffer (Bio-Rad) supplemented with β-mercaptoethanol.

### Immunoblot analysis

Cell lysates or IP eluates were heated for 10 minutes at 95°C prior to resolution on 4-15% SDS-PAGE gradient gels (Bio-Rad), then transferred onto PVDF membranes (Bio-Rad). Membranes were blocked with 5% w/v milk (for Strep Tag II and vinculin antibodies) or 3% w/v bovine serum albumin (for all RNA Pol II antibodies) in Tris-buffered saline and 0.02% Tween 20 (TBST) before incubating with the following primary antibodies at 4°C overnight: RNA Pol II CTD (Clone: 8WG16; Abcam, Ab817; 1:1,000); RNA Pol II CTD phospho-Tyr1 (Clone: 3D12; Active Motif, 61384; 1:1,000); RNA Pol II CTD phospho-Ser2 (Clone: AB_2793780; Active Motif, 91116; 1:1,000); RNA Pol II CTD phospho-Thr4 (Clone: 10A5H5; Invitrogen, MA5-51557; 1:1,000); RNA Pol II CTD phospho-Ser5 (Clone: 4H8; Invitrogen, MA1-46093; 1:1,000); RNA Pol II CTD phospho-Ser7 (Invitrogen PA5-121324,; 1:1,000); Strep Tag II (Clone: 1810CT579.47.56.10; Invitrogen, MA5-37747; 1:500); vinculin (Abcam, 91459; 1:1,000); KSHV ORF34 (homemade; 1:500); KSHV ORF6 sera (1:10,000); KSHV K8.1 (1:10,000). After washing with TBST, membranes were incubated with secondary antibodies at room temperature for 1 hour: Goat Anti-Rabbit IgG(H+L)-HRP (SouthernBiotech, 4050-05; 1:3,000), Goat Anti-Mouse IgG(H+L)-HRP (SouthernBiotech, 1031-05; 1:3,000), or Goat Anti-Rat IgG(H+L)-HRP (SouthernBiotech, 3051-05; 1:3,000). Membranes were incubated with Clarity™ Western ECL Substrate (Bio-Rad) and visualized by chemiluminescence on an Azure 600 imaging system.

### BAC mutagenesis and iSLK establishment

Mutations (N42A, F64A, R68A, R95A) in the ORF24 locus were introduced by Red recombineering of KSHV BAC16 containing an N-terminal 1xHA epitope tag on ORF24 in *Escherichia coli* GS1783 as previous described ^26,39,70,86^. Modified BACs were purified using a NucleoBond Xtra BAC kit (Takara Bio) and digested with RsrII (New England Biolabs) to ensure that no large-scale unintended recombination occurred during cloning.

HEK293T cells constitutively expressing transcomplementing ORF24 for iSLK establishment were generated by lentiviral transduction. Lentivirus was generated in HEK293T cells by cotransfection of pLJM1-HA-ORF24 and packaging plasmids pMD2.G and psPAX2. Two days later, the supernatant was syringe filtered through a 0.45-μm filter. The supernatant was diluted 1:2 with DMEM, and polybrene (Sigma-Aldrich) was added to a final concentration of 8 μg/ml on 1 × 10^6^ freshly trypsinized HEK293T cells and spinoculated in a 6-well plate for 2 hours at 876 × g. After cells had readhered to the plate, the media was replaced with fresh DMEM+10% FBS. The next day, cells were expanded to a 10-cm tissue culture plate and selected for 2 weeks in media supplemented with 500 μg/ml zeocin (Invitrogen).

Latently infected iSLK cell lines were generated via transfection of HEK293T-ORF24 cells with 10 μg BAC DNA using PEI MAX (MW 40,000) transfection reagent (Polysciences, Inc.). The next day, transfected HEK293T cells were trypsinized and mixed 1:1 with iSLK-puro cells and treated with 30 nM 12-O-tetradecanoylphorbyl-13-acetate (TPA) and 0.3 mM sodium butyrate for 4 days to induce lytic replication. iSLK cells were selected for one week in DMEM+FBS containing 300 μg/mL hygromycin B, 1 μg/mL puromycin, and 250 μg/mL G418. The hygromycin B concentration was progressively increased to 1 mg/mL until selection was complete.

### Supernatant transfer to monitor infectious virion production

2.6 × 10^6^ iSLK cells were plated in 15-cm dishes and reactivated into the lytic cycle the next day via addition of 1 μg/ml doxycycline and 1 mM sodium butyrate. 72 hours later, viral supernatant was 0.45 μm filtered and 250 μL of supernatant was spinoculated onto 1 × 10^6^ HEK293T cells for 2 hours at 1,000 × g in a 6-well plate. 24 hours later, the infected HEK293T cells were washed with DPBS and crosslinked in 4% paraformaldehyde (Electron Microscopy Services) in DPBS. The cells were pelleted and resuspended in DPBS and 10,000 cells/sample were analyzed on a BD Accuri 6 flow cytometer. The data were analyzed using FlowJo software (version 10).

### Fluorescent labeling of purified 26xCTD

GST-26xCTD was labeled with Texas Red-X, Succinimidyl Ester, mixed isomers (Invitrogen). Five milligrams of the dye was dissolved in 500 μL of DMSO, and 100 μL of the dissolved dye was incubated with 1 mL of 10 mg/mL GST-26xCTD protein sample. The protein-dye mixture was incubated for 1 hour at 4 °C with constant rotation. The free dye was removed using Micro Bio-Spin P-6 Gel column (Bio-Rad) pre-equilibrated with PBS. The labeled protein was concentrated and buffer exchanged into 150 mM NaCl, 50 mM Tris pH 7.5, 10% glycerol, 1 mM DTT using a 30 K Amicon Ultra-15 concentrator (MilliporeSigma) before aliquoting and storing at -80°C.

#### *In vitro* confocal microscopy

Fluorescent proteins (GFP or GFP-ORF24-NTD-strep) and Texas Red-labeled 26xCTD were incubated with 16% dextran in crowding buffer (150 mM NaCl, 50 mM Tris (pH 7.5), 10% glycerol, 1 mM DTT) at indicated concentrations for 30 minutes. Then, 10 μL of samples were loaded onto glass slides and covered with 22 mm coverslips. Fluorescent images were collected on an LSM 880 laser scanning confocal microscope (Zeiss) with a 100x Plan Apo objective (oil immersion).

#### Turbidity assay

Samples (50 μL) of purified recombinant GST-26xCTD, ORF24-NTD-strep, GFP, or GFP-ORF24-NTD-strep and dextran (MilliporeSigma) in 150 mM NaCl, 50 mM Tris, 10% glycerol, 1 mM DTT, pH 7.5 were prepared at varying concentrations in a clear polystyrene flat bottom microplate 96-well plate (Greiner). Samples were incubated at 30°C in a Synergy MX plate reader (BioTek Instruments) with medium shaking for 5 minutes before measurement of the absorbance at 350 nm.

### Sequence alignment

Closely related gammaherpesvirus homologs of KSHV ORF24 were identified via NCBI BLAST using the KSHV BAC16 ORF24 sequence as a search seed. Sequences with the following accession numbers were aligned in COBALT: ALH45230.1 (KSHV), YP_010801652.1 (Colobine gammaherpesvirus 1; CbGHV1), YP_010084388.1 (Retroperitoneal fibromatosis-associated herpesvirus; RFHV), NP_570764.1 (Macacine gammaherpesvirus 5; McHV-5), YP_010084566.1 (Macaca nemestrina rhadinovirus 2; McRV2), NP_076516.1 (Bovine gammaherpesvirus 4; BoHV-4), NP_047995.1 (Ateline gammaherpesvirus 3; AtHV-3), NP_040226.1 (Saimiriine gammaherpesvirus 2; SaHV-2), YP_009173899.1 (Felid gammaherpesvirus 1; FeHV-1), YP_009229858.1 (Myotis gammaherpesvirus 8; BGHV8), UNP64590.1 (Marmot herpesvirus 1; MHV-1), and UNP64498.1 (Saguinine gammaherpesvirus 1; SgHV-1). Sequences were visualized in JalView and colored based on percentage of sequence identity, where >80% is colored dark blue, >60% colored light blue, >40% colored light grey, and <40% colored white.

## Data availability

Atomic coordinates and structure factors for ORF24-NTD have been deposited in the Protein Data Bank with accession code 13DH. Diffraction images have been deposited in the SBGrid Data Bank under ID 1267.

## ACKNOWLEDGEMENTS

We thank Y. Jessie Zhang for 26xCTD-T4E, S5E, and S7E and DYRK1a and ERK2 kinase plasmids and Karla Neugebauer for phosphorylation state-specific anti-CTD antibodies. We thank James Murphy from the Macromolecular X-ray Crystallography Facility at Yale School of Medicine, the Ailong Ke lab, and Rohini Dwivedi for assistance with protein crystallization and data collection. Mass spectrometry data collection and analysis was performed by Jeremy Balsbaugh and Jennifer Liddle at the University of Connecticut Proteomics & Metabolomics Facility. Use of the Orbitrap Eclipse Tribrid mass spectrometer housed in the University of Connecticut Proteomics & Metabolomics Facility was supported by NIH S10 High-End Instrumentation Award 1S10-OD028445-01A1. We thank colleagues at SSRL BL12-2 for assistance with remote data collection. The SSRL Structural Molecular Biology Program is supported by the DOE Office of Biological and Environmental Research, and by the National Institutes of Health, National Institute of General Medical Sciences (P30GM133894).

We thank all members of the Didychuk Lab as well as Leah Gulyas, Mandy Muller, Dan DiMaio, and Britt Glaunsinger for feedback. This work was supported by an NIH DP2 AI171113 award and Damon Runyon Cancer Research Foundation Dale F. Frey Award for Breakthrough Scientists awarded to A.L.D. T.G. was supported by the NSF Graduate Research Fellowship Program (DGE-2139841) and S.T. was supported by an NIH NICHD grant (R00HD104924).

## SUPPLEMENTARY FIGURES

**Supplementary Figure S1.**
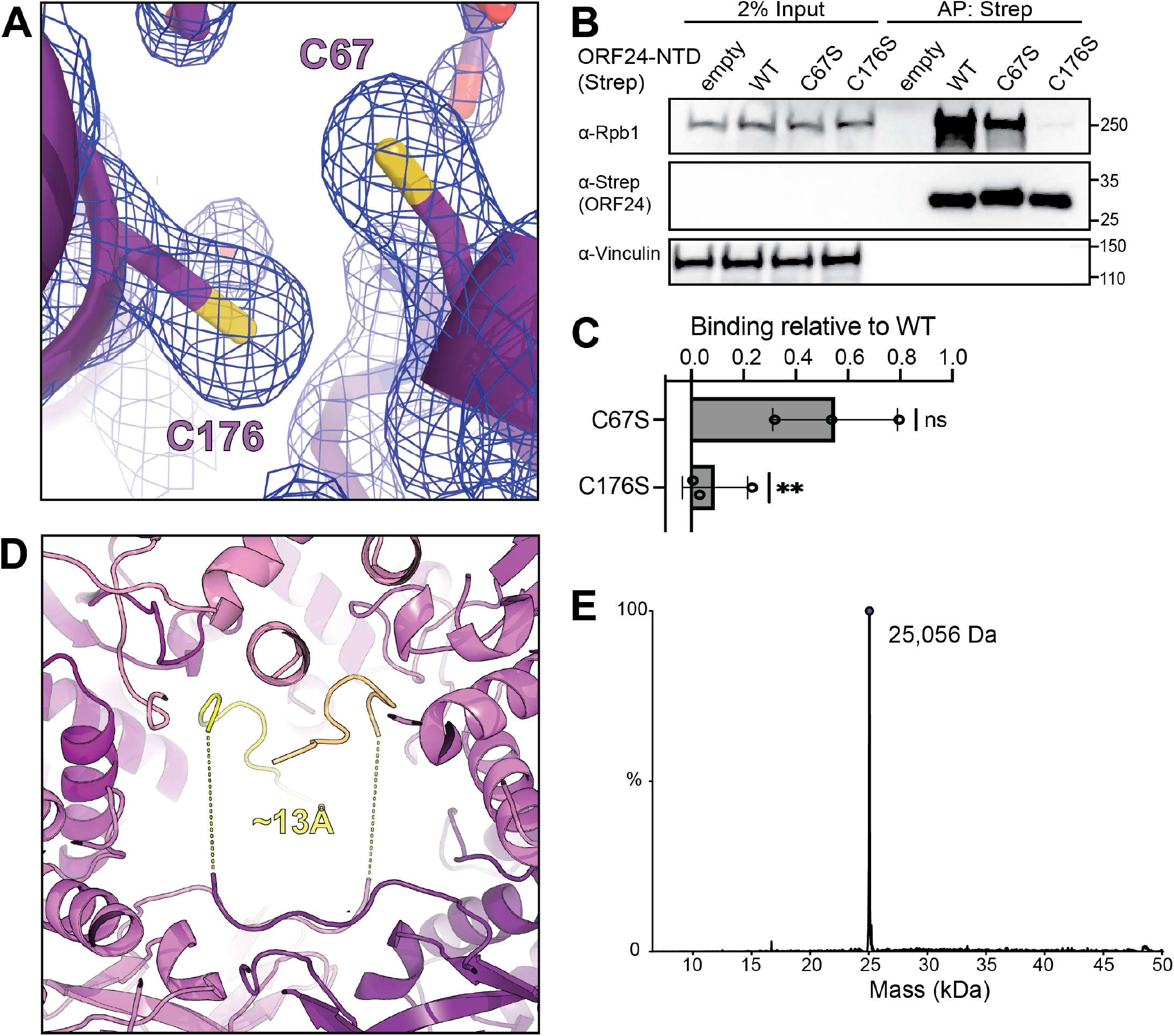
Structural features of ORF24-NTD. **(A)** *2mFo-DFc* electron density map of ORF24-NTD residues C67 and C176, contoured at 1σ. **(B)** Plasmids encoding wild-type or mutant strep-tagged ORF24-NTD were transiently transfected into HEK293T cells and subjected to affinity purification on StrepTactinXT beads followed by western blot analysis; representative of three biological replicates. Molecular weight (in kDa) was determined via a protein ladder and is indicated to the right. Vinculin serves as a loading control. **(C)** Quantification of three independent replicates of the affinity purification experiment shown in (B). The ratio of Rpb1 to ORF24-NTD was normalized to WT in each experiment. Data are from three biological replicates, with statistics calculated using a one sample t-test where WT binding was set to 1; \*\**P* < 0.01. **(D)** The N-terminal hexahistidine tag may mediate crystal packing. The C-terminal ends of the modeled tag (SHHHHHHSSE) from two asymmetric units (yellow, wheat) are ∼13 Å from the N-terminal (-1) Glycine. The intervening sequence (NLYFQ) is likely unstructured and spans this distance. ORF24-NTD is shown in shades of purple. **(E)** UPLC-MS/MS analysis of recombinantly purified His-ORF24-NTD was performed using a C4 BEH analytical column and reversed-phase gradient elution. The MS1 data was averaged across the 54 min elution profile and deconvoluted using UniDec to yield an average molecular weight measurement. The observed MS1 data correlates with an intact average mass of 25,056 Da, which closely matches the expected average molecular weight of unmodified His-ORF24-NTD minus the initiating Met (-131 Da). No other abundant ion signals were observed.

**Supplementary Figure S2.**
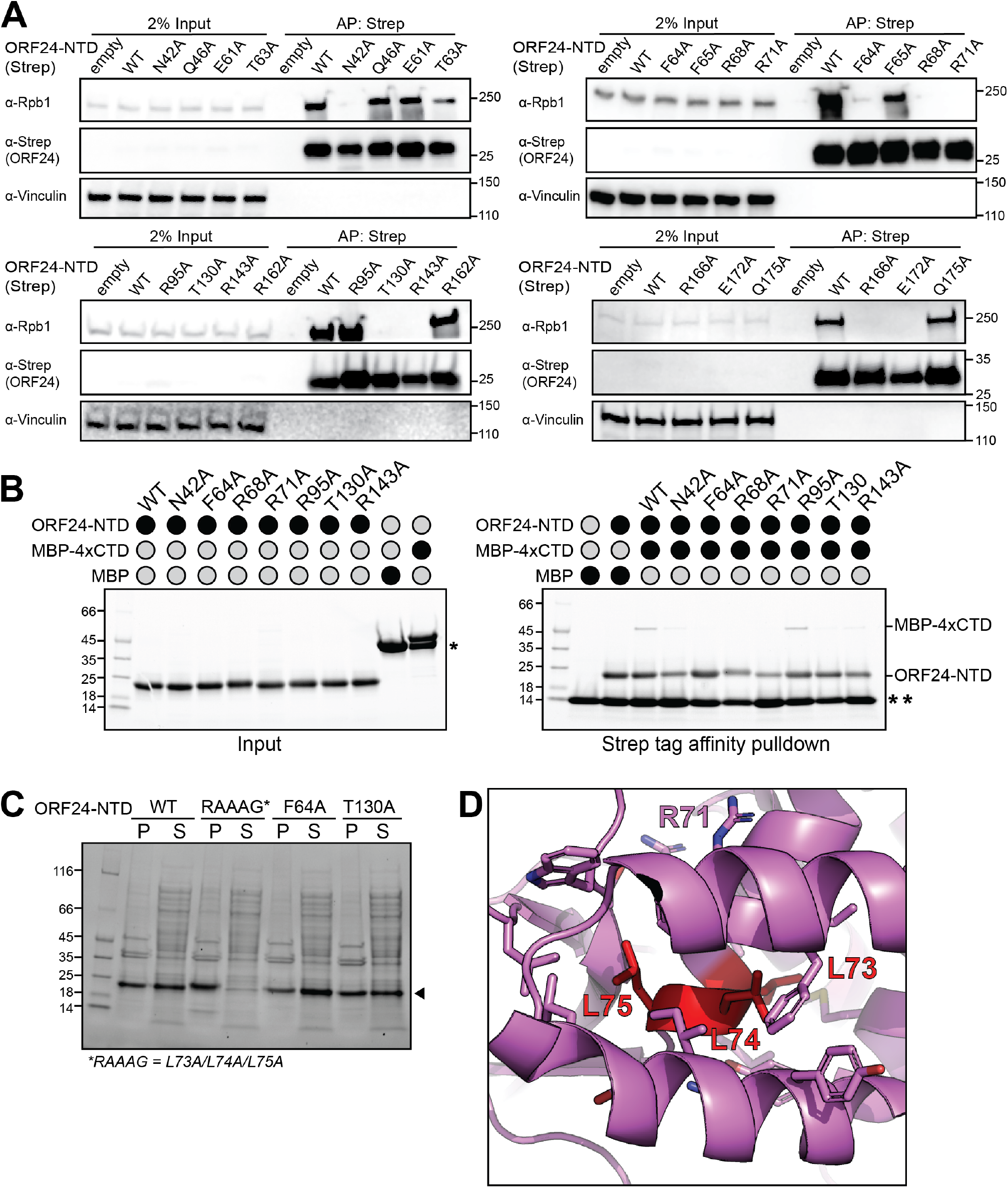
Binding assays to monitor ORF24-NTD/RNAPII CTD interactions. **(A)** Plasmids encoding wild-type or mutant strep-tagged ORF24-NTD were transiently transfected into HEK293T cells and subjected to affinity purification on StrepTactinXT beads followed by western blot analysis. Data are representative of three biological replicates. Molecular weight (in kDa) was determined via a protein ladder and is indicated to the right. Vinculin serves as a loading control. Data are quantified in **Figure 2B. (B)** Representative *in vitro* binding assay where recombinantly purified wild-type or mutant His-ORF24-NTD-strep was incubated with MBP-4xCTD followed by enrichment on StrepTactin XT beads. Eluted fractions were analyzed by stain-free SDS-PAGE. (*) indicates free MBP. (**) indicates a subunit of Strep-TactinXT released from the beads during boil elution. Data are quantified in **Figure 2D. (C)** Solubility analysis of His-ORF24-NTD mutants. Mutants were expressed in *E. coli* under identical conditions. After induction, equal amounts of cells were lysed, and the soluble and insoluble fractions were analyzed by stain-free SDS-PAGE. **(D)** Location of the “RAAAG” (L73A/L74A/L75A) mutation (red) on the ORF24-NTD structure (purple). These residues are buried in the interior of the protein between the base of the β-hairpin and α-helices α1 and α4-6.

**Supplementary Figure S3.**
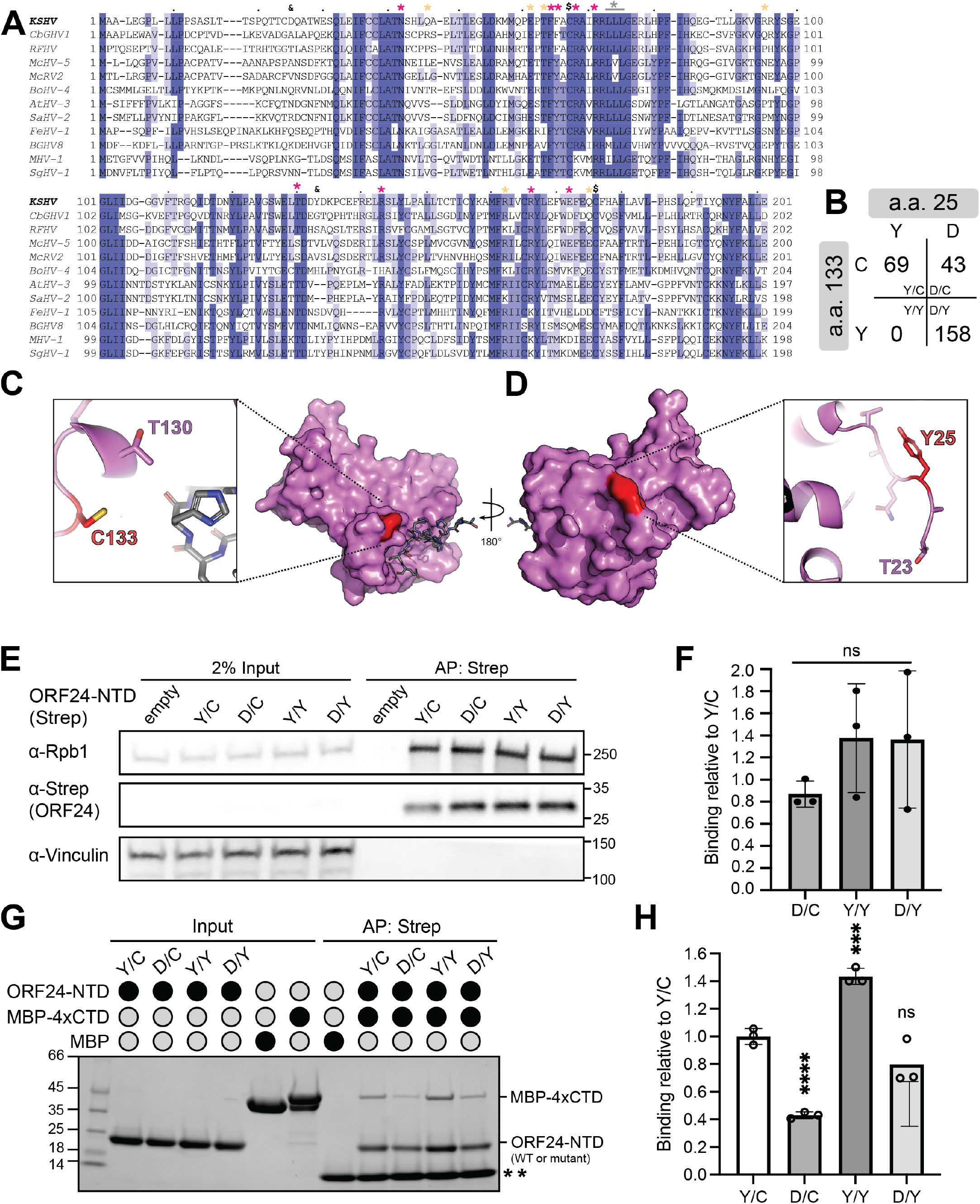
Natural variation at ORF24 residues 25 and 133. **(A)** Multiple sequence alignment of closely related gammaherpesvirus homologs of ORF24 where every tenth residue is indicated with a period above the alignment. Residues critical (pink) or not required for binding (yellow) as determined in Fig. 2 are indicated with asterisks. The grey asterisk marks site of the RAAAG mutations. Dollar signs indicate conserved cysteine residues with the potential to form a disulfide bond. Ampersands indicate positions 25 and 133 that are variable within sequenced KSHV genomes. **(B)** Distribution of naturally occurring residues present at positions 25 and 133 from 270 sequenced KSHV genomes available in NCBI. **(C)** Position and local contacts for residues C133 and **(D)** Y25 in the ORF24-NTD structure, highlighted in red sticks Nearby residues are shown in purple sticks, with the putative hexahistidine peptide shown in grey sticks. **(E)** Plasmids encoding ORF24-NTD harboring different combinations of residues at positions 25 and 133 were transiently transfected into HEK293T cells and subjected to affinity purification on StrepTactinXT beads followed by western blot analysis; representative of three biological replicates. Molecular weight (in kDa) was determined via a protein ladder and is indicated to the right. Vinculin serves as a loading control. **(F)** Quantification of three independent replicates of the affinity purification experiment shown in (E). The ratio of Rpb1 to ORF24-NTD was normalized to WT in each experiment. Data are from three biological replicates, with statistics calculated using a one sample t-test where WT binding was set to 1. **(G)** *In vitro* binding assay where recombinantly purified MBP-4xCTD and variants of His-ORF24-NTD-strep were enriched on StrepTactin XT beads. Free MBP was used as a negative control. (**) indicates a subunit of Strep-TactinXT released from the beads during boil elution. **(H)** Quantification of three independent replicates of the *in vitro* binding assay shown in (G). Data from three independent replicates were normalized to the average of wild-type binding, with statistics calculated using an unpaired t-test; \*\*\*\**P* < 0.0001, \*\*\**P* < 0.001.

**Supplementary Figure S4.**
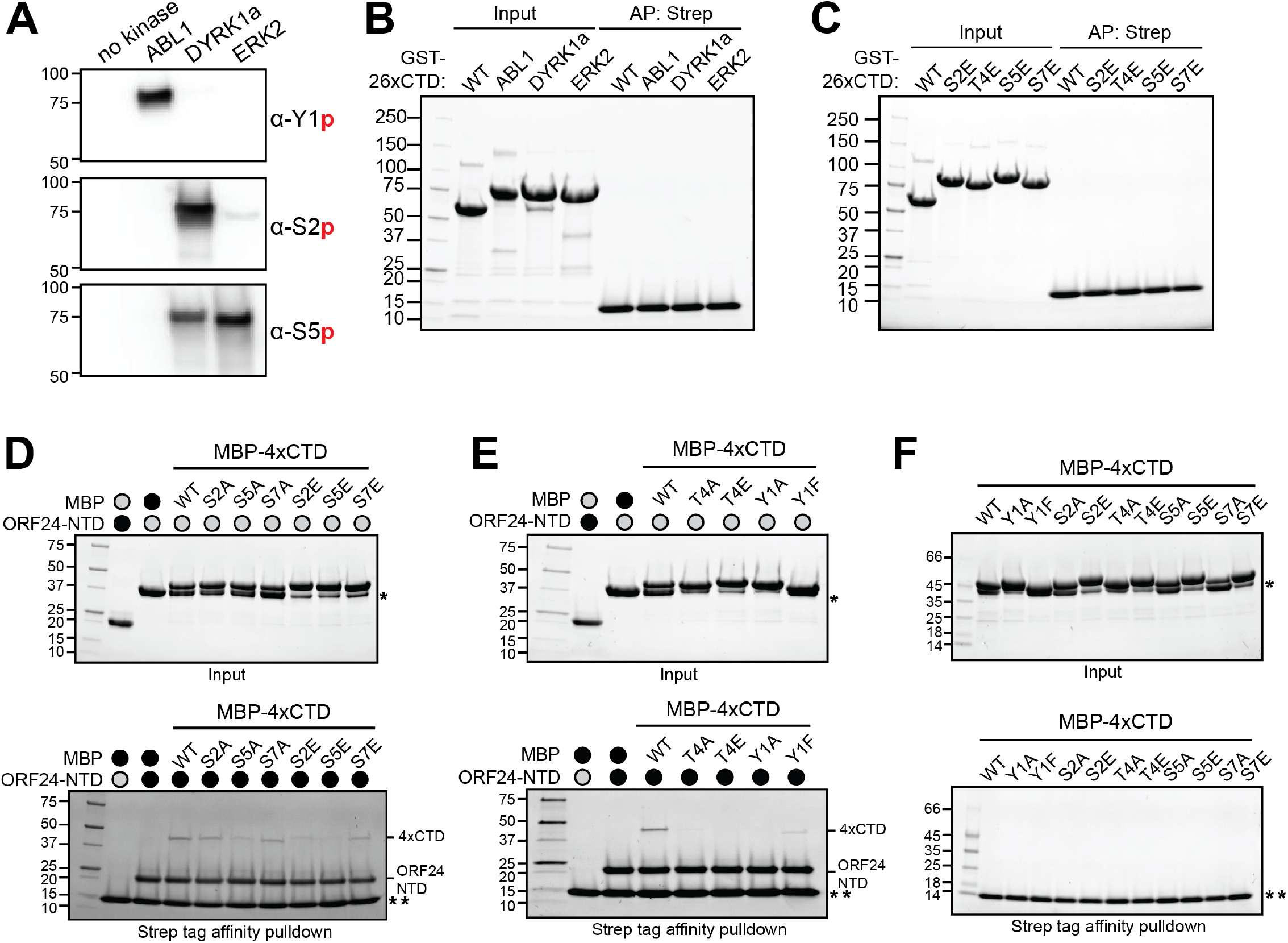
Supporting data for phosphorylation of RNAPII CTD and mutant binding data. **(A)** Phosphorylated GST-26xCTD (0.2 μg) was analyzed by western blot with phospho-specific antibodies. **(B)** Phosphorylated GST-26xCTD or **(C)** GST-26xCTD phosphomimetic variants are not enriched by Strep-TactinXT beads in the absence of His-ORF24-NTD-strep. **(D)** Positions S2, S5, S7 and **(E)** Y1, T4 in the CTD heptad sequence affect binding of ORF24-NTD. Representative gels of an *in vitro* binding experiment quantified in Figure 4 using MBP-4xCTD mutants binding to His-ORF24-NTD-Strep. The top gel shows input (wild-type and mutant MBP-4xCTD and His-ORF24-NTD-strep) while the bottom gel shows the resulting enrichment from StrepTactinXT beads. (*) indicates free MBP, (**) indicates a subunit of Strep-TactinXT released from the beads during boil elution. Gels were visualized by stain-free imaging. **(F)** MBP-4xCTD variants do not bind StrepTactinXT beads in the absence of His-ORF24-NTD-strep. The top gel shows input (free MBP-4xCTD variants) and the bottom gel shows the resulting enrichment from StrepTactinXT beads.

**Supplementary Figure S5.**
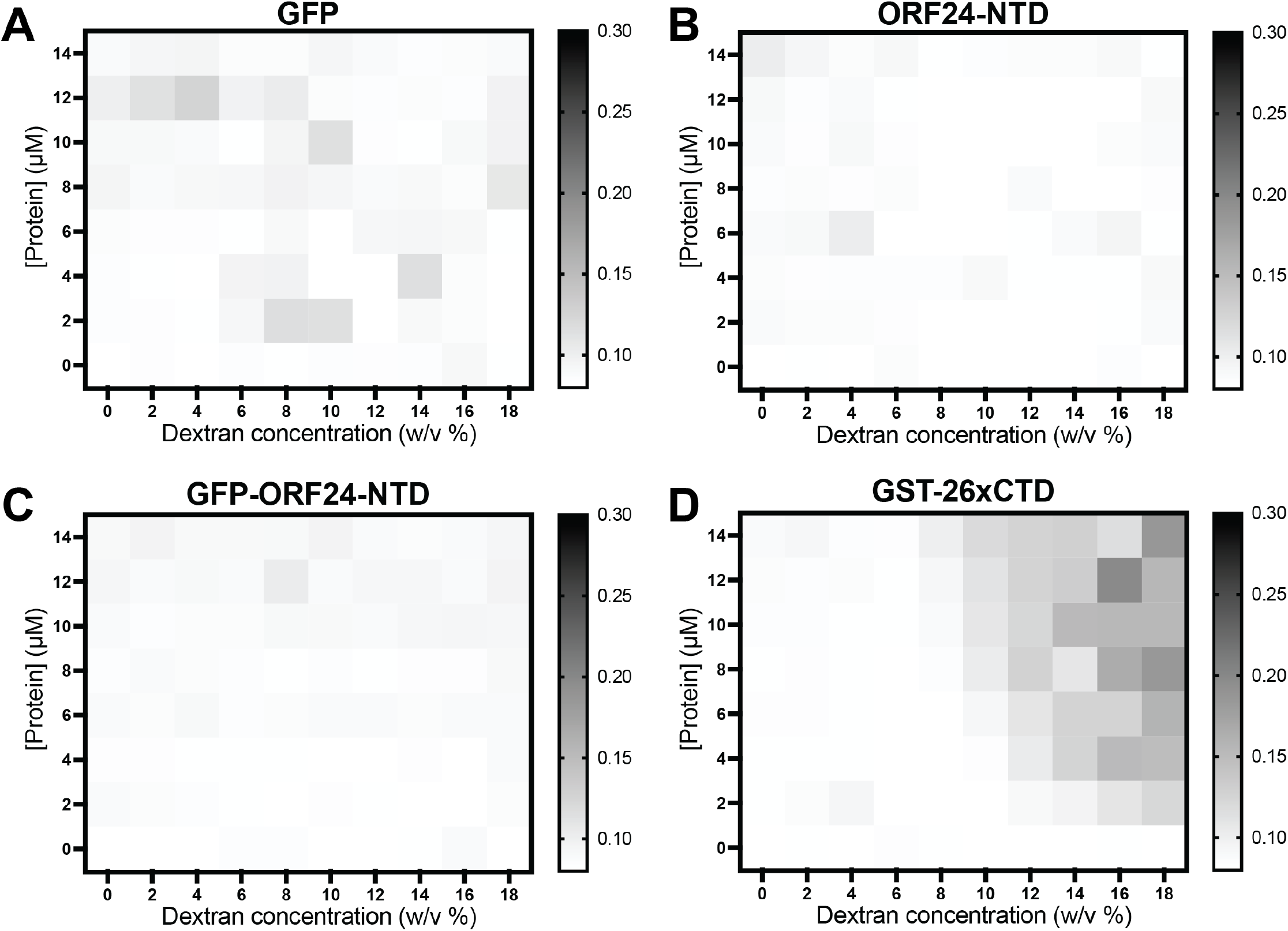
RNAPII CTD, but not ORF24, phase separates *in vitro*. Condensate formation at various concentrations of protein and dextran was monitored via absorbance at 350 nm for **(A)** free GFP, **(B)** ORF24-NTD-strep, **(C)** GFP-ORF24-NTD-strep, or **(D)** GST-26xCTD. Graphs represent the average of two independent replicates.

**Supplementary Table 1.** Crystallographic data collection, phasing and structure refinement statistics.

| Parameter | ORF24-NTD (PDB 13DH) |
| --- | --- |
| **Data Collection** |  |
| Wavelength (Å) | 0.9795 |
| Resolution range (Å) | 36.75 - 1.932 (2.08 - 1.93) |
| Space group | P 41 21 2 |
| Unit cell |  |
| a, b, c (Å) | 51.97, 51.97, 147.6 |
| $\alpha=\beta=\gamma$ (°) | 90 |
| Total reflections | 227,717 |
| Unique reflections | 13,138 (1,143) |
| Multiplicity | 17.3 |
| Completeness (%) | 82.39 (36.82) |
| Mean I/sigma(I) | 17.8 |
| Wilson B-factor | 38.86 |
| R-merge | 10.50% |
| R-meas | 10.80% |
| CC1/2 | 12.1 |
| CC* | 99.9 |
| **Refinement Statistics** |  |
| Resolution range (Å) | 36.75 - 2.04 (2.04 - 1.93) |
| R-work / R-free | 0.1860 / 0.2434 (0.3474/0.3684) |
| Number of non-hydrogen atoms | 1891 |
| macromolecules | 1713 |
| ligands | 60 |
| solvent | 118 |
| Protein residues | 213 |
| RMS(bonds) | 0.011 |
| RMS(angles) | 1.76 |
| Ramachandran favored (%) | 96.17 |
| Ramachandran allowed (%) | 3.35 |
| Ramachandran outliers (%) | 0.48 |
| Rotamer outliers (%) | 6.45 |
| Clashscore | 3.42 |
| Average B-factor | 58.11 |
| macromolecules | 56.95 |
| ligands | 87.36 |
| solvent | 60 |
| Number of TLS groups | 2 (for protein and peptide) |

